# Structure-function analyses of dual-BON domain protein DolP identifies phospholipid binding as a new mechanism for protein localisation

**DOI:** 10.1101/2020.08.10.244616

**Authors:** J. A. Bryant, F. C. Morris, T. J. Knowles, R. Maderbocus, E. Heinz, G. Boelter, D. Alodaini, A. Colyer, P. J. Wotherspoon, K. A. Staunton, M. Jeeves, D. F. Browning, Y. R. Sevastsyanovich, T. J. Wells, A. E. Rossiter, V. N. Bavro, P. Sridhar, D. G. Ward, Z-S. Chong, C. Icke, A. Teo, S-S. Chng, D. I. Roper, T. Lithgow, A. F. Cunningham, M. Banzhaf, M. Overduin, I. R. Henderson

## Abstract

The Gram-negative outer membrane envelops the bacterium and functions as a permeability barrier against antibiotics, detergents and environmental stresses. Some virulence factors serve to maintain the integrity of the outer membrane, including DolP (formerly YraP) a protein of unresolved structure and function. Here we reveal DolP is a lipoprotein functionally conserved among Gram-negative bacteria and that loss of DolP increases membrane fluidity. We present the NMR solution structure for DolP, which is composed of two BON domains that form an interconnected opposing pair. The C-terminal BON domain binds to anionic phospholipids through an extensive membrane:protein interface providing evidence of subcellular localization of these phospholipids within the outer membrane. This interaction is essential for DolP function and is required for sub-cellular localization of the protein to the cell division site. The structure of DolP provides a new target for developing therapies that disrupt the integrity of the bacterial cell envelope.

## Main

Gram-negative bacteria are intrinsically resistant to many antibiotics and environmental insults, which is largely due to the presence of their hydrophobic outer membrane (OM). This asymmetric bilayer shields the periplasmic space, a thin layer of peptidoglycan and the inner membrane (IM). In the model bacterium *Escherichia coli*, the IM is a symmetrical phospholipid bilayer, whereas the OM has a more complex organisation with lipopolysaccharide (LPS) and phospholipids forming an asymmetric bilayer containing integral β-barrel proteins ^1,2^. The OM is also decorated with lipoproteins (approximately 75 have been identified in *E. coli*), many of which, are functional orphans ^3,4^. Biogenesis of the OM is completed by several proteinaceous systems, which must bypass the periplasmic, mesh-like peptidoglycan ^2,5–7^. The growth of all three envelope layers must be tightly co-ordinated in order to maintain membrane integrity. Improper coordination can lead to bacterial growth defects, sensitivity to antibiotics, and can cause cell lysis ^5,8^.

DolP (**<u>d</u>**ivision and **<u>O</u>**M stress-associated **<u>l</u>**ipid-binding **<u>p</u>**rotein; formerly YraP) is a nonessential protein found in *E. coli* and other Gram-negative bacteria ^9^. Loss of DolP results in the disruption of OM integrity, induces increased susceptibility to detergents and antibiotics, and attenuates the virulence of *Salmonella enterica* ^10^. Importantly, DolP is a crucial component of the serogroup B meningococcal vaccine where it enhances the immunogenicity of other components by an unknown mechanism ^11^. Recently, the *dolP* gene was connected genetically to the activation of peptidoglycan amidases during *E. coli* cell division, however this activity has not been directly confirmed experimentally ^12^. In contrast, protein interactome studies suggest DolP is a component of the β-barrel assembly machine (Bam) complex ^13,14^. While these data suggest that DolP may be involved in outer membrane protein (OMP) biogenesis and the regulation of peptidoglycan remodeling, its precise function in either of these processes remained unclear. Nonetheless, given its roles in these vital cell envelope processes, and its requirement for virulence and the maintenance of cell envelope integrity, DolP is a potential target for the development of therapeutics.

In this study, we demonstrate that DolP is an outer membrane lipoprotein functionally conserved amongst Gram-negative bacteria, but with a function distinct from other BON domain containing proteins. We solve the NMR solution structure of DolP revealing the first view of a dual-BON domain fold. Extensive structural and functional analyses define a membrane:protein interface that binds DolP to anionic phospholipids and provides the basis for a new mechanism for targeting proteins to the cell division site. We show that loss of *dolP* affects OM fluidity, which perturbs the BAM complex, suggesting an indirect role for DolP in OMP biogenesis. The insights provided here not only advance our understanding of how DolP functions but provide a basis for beginning to develop drugs to target it.

## Results

### DolP belongs to an extensive family of lipoproteins required for OM homeostasis

In *E. coli*, the *dolP* gene is located downstream of the genes encoding LpoA (an activator of PBP1A) ^15^, YraN (a putative Holiday-Junction resolvase), and DiaA (a regulator of chromosomal replication) ^16^, and two σ^E^-dependent promoters are found immediately upstream of the *dolP* gene ^17^ **(Fig. 1A)**. Bioinformatic analyses predicted that *dolP* encodes a lipoprotein with two putative domains of unknown function, termed BON domains ^18^, as well as a Lol-dependent OM targeting signal sequence where acylation was predicted to occur on cysteine residue C19. To test the hypothesis that DolP is localized to the periplasmic face of the OM, we raised an antiserum to the protein to probe subcellular fractions. DolP was found in the Triton X-100 insoluble fraction of the *E. coli* cell envelope along with other OM proteins. As a control for the antiserum, DolP was absent from Triton X-100 insoluble fractions of cell envelopes harvested from *E. coli* Δ*dolP* **(Fig. S1A)**. Furthermore, expression of a C19A point mutant, preventing N-terminal acylation, effectively eliminated DolP from the OM fractions **(Fig. S1B).** Unlike the lipoproteins BamC and Lpp, which can be surface localised ^19,20^, DolP was not accessible to antibody or protease in intact *E. coli* cells. However, DolP could be labelled and degraded when OM integrity was compromised **(Fig, S1, C and D)**, confirming that DolP is predominantly targeted to the inner leaflet of the OM, localizing it within the periplasmic space.

**Fig. 1.**
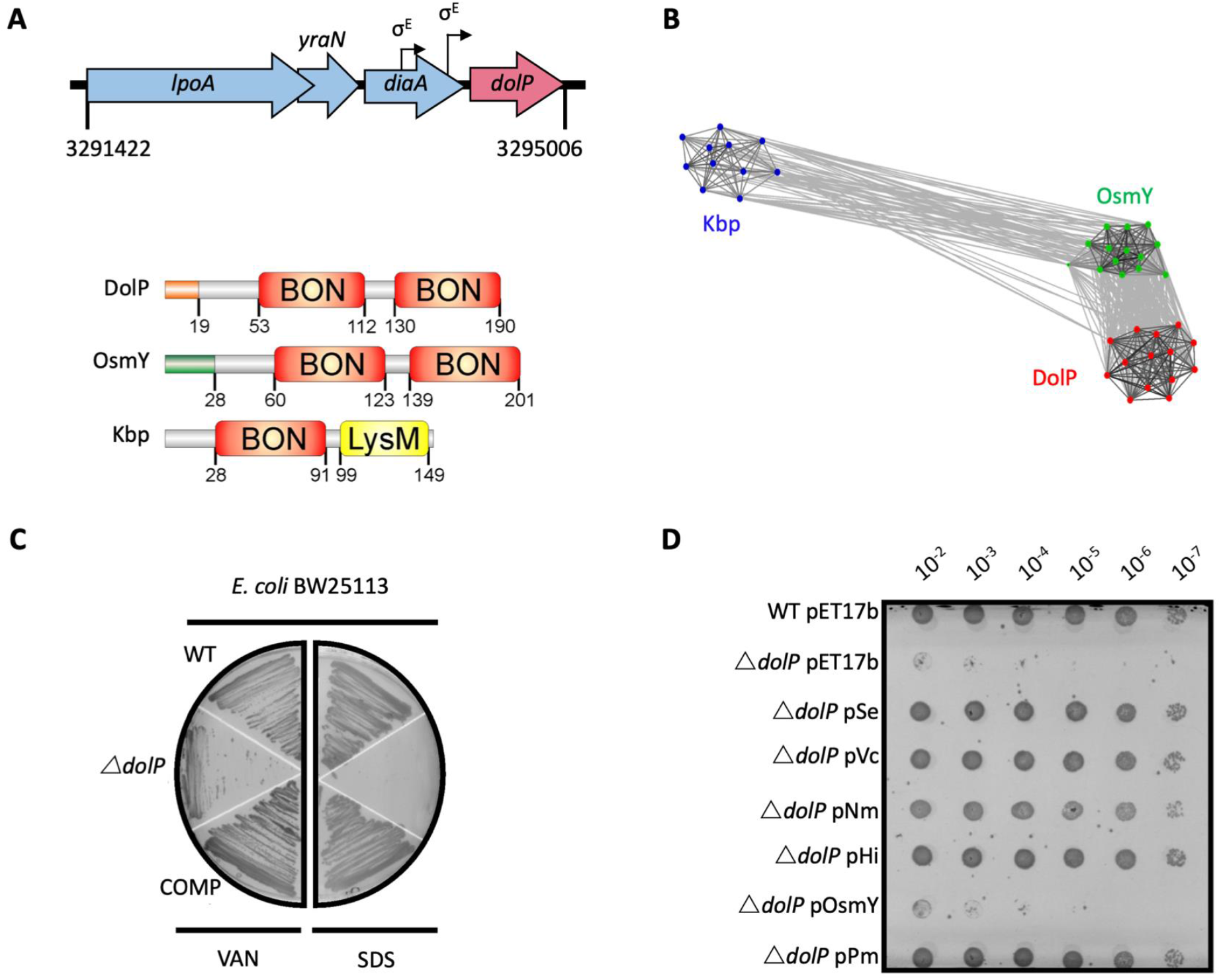
DolP is a conserved BON domain protein with a distinct role in OM homeostasis. **A.** In *E. coli*,*dolP* is located downstream of *diaA* and encodes a lipoprotein with a signal sequence (orange) and two BON domains (red). The signal sequence is cleaved by LspA, the cysteine at position 19 acylated by Lgt and Lnt and finally the protein is targeted to the OM by the Lol system **(Fig. S1)**. *E. coli* contains three BON domain proteins. DolP shares a similar domain organisation with OsmY, which encodes a periplasmic protein that possesses a signal sequence (green) which is recognised and cleaved by the signal peptidase LepB. Kbp is more divergent from DolP and OsmY, has no predictable signal sequence and is composed of BON and LysM domains **(Fig. S2)**. **B.** DolP, OsmY and Kbp are widespread among proteobacteria, and cluster into three distinct groups based on the program CLANS43 with connections shown for a *P* value cut-off of <10^−2^ **(Table S5)**. **C.** Growth phenotypes for mutant isolates lacking DolP (Δ *dolP*), wild-type strain (WT) or the complemented mutant (COMP). Strains were grown on LB agar containing vancomycin (100 μg/ml) or sodium dodecyl sulphate (SDS; 4.8%). **D.** DolP from diverse proteobacterial species expressed in an *E. coli* Δ*dolP* strain restores growth in the presence of vancomycin as assessed by a serial dilution plate growth assay. Plasmids expressing OsmY do not complement the defect.

Further in silico analyses revealed the DolP lipoprotein was conserved across diverse species of Proteobacteria and is present even in organisms with highly-reduced genomes e.g. *Buchnera* spp **(Tables S1 and S2).** The genome of *E. coli* contains three BON domain-containing proteins: DolP, OsmY and Kbp. DolP shares a dual BON-domain architecture and 29.5% sequence identity with OsmY, which is distinguished from DolP by a canonical Sec-dependent signal sequence. In contrast, Kbp consists of single BON and LysM domains and lacks a discernible signal sequence **(Fig. 1A)**. Our comprehensive analysis found seven predominant domains co-occurring with BON in different modular protein architectures across bacterial phyla, suggesting specialized roles for BON domains **(Table S1 and Fig. S2)**. Clustering analyses of sequences obtained by HMMER searches revealed DolP, OsmY and Kbp are distributed throughout the α, β and γ-proteobacteria and form distinct clusters indicating that DolP has a role that is independent of OsmY and Kbp **(Fig. 1B).** Our analyses demonstrated that OsmY and Kbp are not functionally redundant with DolP and isogenic mutants show distinct phenotypes, therefore confirming a distinct role for DolP in *E. coli* **(Fig. S3).**

Previously, we demonstrated that loss of *dolP* in *S. enterica* conferred susceptibility to vancomycin and SDS, suggesting DolP plays an important role in maintaining the integrity of the OM ^10^. Further evidence of a role for DolP in maintaining OM integrity is shown by *E. coli* Δ*dolP* susceptibility to vancomycin, SDS, cholate, and deoxycholate **(Fig. 1C and Fig. S4A).** Resistance could be restored by supplying *dolP* in *trans* **(Fig. 1C)**. Despite evidence for disrupted OM integrity, the growth rate observed for the *dolP* mutant strain was identical to that of the parent, and scanning-electron microscopy revealed no obvious differences in cell size or shape **(Fig. S4, B and C)**. To determine whether DolP is broadly required for OM homeostasis, plasmids expressing DolP homologues from *S. enterica*, *Vibrio cholerae*, *Pasteurella multocida*, *Haemophilus influenza* and *Neisseria meningitidis* were shown to restore the OM barrier function of the *E. coli ΔdolP* mutant **(Fig. 1F)**. Finally, either replacement of the DolP signal sequence with that of PelB, which targets the protein to the periplasmic space, or mutation of the signal sequence to avoid OM targeting *via* the Lol system, prevented complementation of the Δ*dolP* phenotype **(Fig. S5)**. Together these results support a conserved role for DolP in maintenance of OM integrity throughout Gram-negative bacteria and demonstrate that localization of DolP to the inner leaflet of the OM is essential to mediate this function.

### The structure of DolP reveals a dual-BON domain lipoprotein

To gain further insight into the function of DolP, the structure of full-length mature *E. coli* DolP was determined by NMR spectroscopy. To promote native folding of DolP, the protein was over-expressed in the periplasm using a PelB signal sequence; the N-terminal cysteine was removed to prevent acylation and provide for rapid purification of the soluble protein. Purified DolP was processed, soluble and monomeric, as confirmed by analytical ultra-centrifugation and size exclusion chromatography **(Fig. S6, A and B)**. Using a standard Nuclear Overhauser Effect (NOE)-based approach, a convergent ensemble was calculated from the ^20^ lowest-energy solution structures, revealing two BON domains facing away from each other and offset by ~45° **(Fig. 2A and Fig. S7).** The individual BON1 and BON2 domains have C-alpha backbone root mean square deviations (RMSDs) of 0.60 and 0.78 Å, respectively, and an overall global RMSD of 1.21 Å **(Table S3)**. Despite having low sequence identity (24.7%) each BON domain consists of a three-stranded mixed parallel/antiparallel β-sheet packed against two α-helices yielding an αββαβ topology. The two BON domains present high structural homology and superpose with an RMSD of 1.8 Å over C-alpha backbone **(Figs. S7 and S8).** Notably, BON1 is embellished by an additional short α1* helix between BON1:α1 and BON1:β1 **(Fig. 2A, Fig. S7 and S8)**. The N-terminal acylation site is connected through a ^27^ amino acid dynamic unstructured linker **(Fig. 2B)**. The molecular envelope of full length DolP calculated by small angle X-ray scattering (SAXS) accommodated the NMR derived structure of DolP and supported the presence of a flexible N-terminal extension. The experimentally determined scattering curve fit the NMR derived structure with a χ^2^ of 1.26, confirming the accuracy of the NMR-derived structure and an exclusively monomeric state **(Fig. 2C)**.

**Fig. 2.**
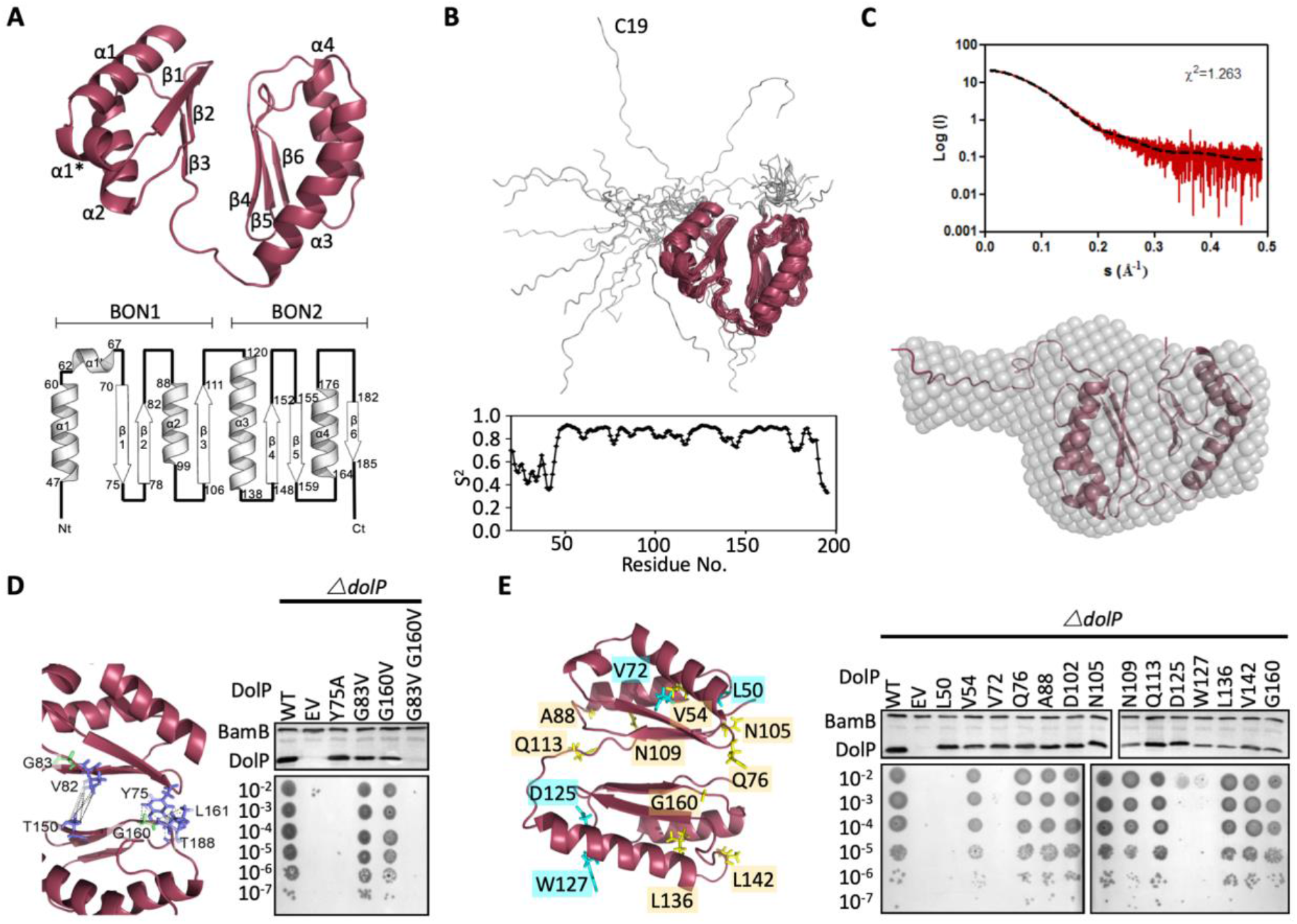
Structure of DolP. **A.** Solution structure and topology of DolP, with α helices, β strands and termini labelled. **B.** Backbone model of the 20 lowest-energy solution structures of DolP. The core folded domain is highlighted in red whilst the flexible N-terminal is shown in grey. The dynamic nature of the linker was demonstrated from S2 order parameter analysis calculated from chemical shift assignments using TALOS+. **C.** Small Angle X-ray Scattering curve of DolP with corresponding best fit of the solution structure of DolP. Best fit calculated based on the core DolP solution structure with flexibility accommodated in residues 20-46, 112-118 and 189-195. The corresponding *ab initio* bead model is shown calculated using Dammif72 based solely on the scattering data. **D.** Structure of DolP showing the conserved residues forming interdomain contacts. Western blots of total protein extracts show plasmid-mediated expression of DolP in *E. coli* Δ*dolP* after site-directed mutation of conserved residues. The empty vector (EV) control is labelled and WT represents wild type DolP. The presence of the OM lipoprotein BamB was used as a control. Colony growth assays by serial dilution of mutants reveal which residues are critical for the maintenance of the OM barrier function. **E.** Structure of DolP showing position of transposon-mediated insertions. Western blots of total protein extracts show plasmid-mediated expression of mutant versions of DolP in *E. coli* Δ*dolP*. The empty vector (EV) control is labelled and WT represents wild type DolP. Colony growth assays by serial dilution of mutants reveal which insertions abolish DolP function. Blue labels represent position of non-functional insertions. Orange labels represent position of tolerated insertions. The presence of the OM lipoprotein BamB was used as a control.

The two BON domains pack against each other *via* their β-sheets through contacts mediated directly by Y75 and V82 in BON1 and T150, G160, L161 and T188 in BON2 **(Fig. 2D)**. This interdomain orientation is consistent with SAXS analysis **(Fig. 2C)** and appears to be essential for function as the mutation Y75A abolishes function **(Fig. 2D).** Single point mutations (G83V and G160V) of the highly conserved glycine residues had less effect, however the double mutant was non-functional **(Fig. 2D and Fig. S8)**. Since the latter protein was not detectable by Western immunoblotting this is likely due to structural instability **(Fig. 2D)**.

The elements of DolP that are required for function were mapped using an unbiased linker-scanning mutagenesis screen. The resulting DolP derivatives, containing in-frame 5-amino-acid insertions, were tested for stability by Western immunoblotting. Functional viability was assessed by their capacity to restore growth of *E. coli* Δ*dolP* in the presence of SDS **(Fig. 2E).** Seven mutants occurred in the signal sequence and the linker region and were not considered further. Eight insertions were identified in BON1, with insertions at positions L50 (BON1:α1) and V72 (BON1:β1) failing to complement the ΔdolP defect whereas the rest were well tolerated. Five insertions were found in BON2, with those at positions L136, L142 and G160 being well tolerated. The remaining insertions at positions D125 and W127 occurred in BON2:α1 but failed to complement the ΔdolP phenotype. None of these mutations abolished protein expression. These data indicate the importance of BON2:α1 in maintaining DolP function and OM integrity **(Fig. 2E)**.

### DolP binds specifically to anionic phospholipids via BON2

Given that OM permeability defects are often associated with the loss or modification of molecular partners, we sought to identify DolP ligands. Published high-throughput protein:protein interaction data^13,14^ suggested that DolP co-located with components of the BAM complex in the OM. Indeed, we observed that simultaneous deletion of *dolP* and genes coding the non-essential BAM complex components *bamB* or *bamE* lead to negative genetic interactions and increased rates of cell lysis **(Fig. S9A and B)**, suggesting a potential interaction. However, despite the genetic interactions, in our hands no significant interaction could be detected between DolP and the BAM complex through immunoprecipitations **(Fig. S9C)** and no significant change in overall OMP levels was observed **(Table S4 and Fig. S9D)**. Analyses of purified OM fractions revealed no apparent differences in LPS profiles **(Fig. S10A)**, or phospholipid content **(Fig. S10B)** between the parent and the *dolP* mutant. No significant increase in hepta-acylated Lipid A was observed in the absence of DolP, indicating that the permeability defect is also not due to loss of OM lipid asymmetry **(Fig. S10C)**. In contrast, Δ*dolP* cells were found to have an increase in membrane fluidity **(Fig. S10D)** as assessed by staining with the membrane intercalating dye pyrene-decanoic acid (PDA), which undergoes a fluorescence shift upon formation of the excimer, an event which is directly related to membrane fluidity^21^. These data suggest that the genetic interaction between *dolP* and *bamE* or *bamB*, observed here, is likely facilitated indirectly through changes to membrane fluidity on the loss of DolP.

Considering that the *dolP* mutant has changes to membrane fluidity and that BON domains are suggested to bind phospholipids^18^, we sought to test whether DolP interacts with phospholipids. A set of potential ligands were screened by chemical shift perturbation (CSP) analysis, including *E. coli* OM lipids embedded in micelles. DolP bound specifically to micelles containing the anionic phospholipids phosphatidylglycerol (PG) and cardiolipin (CL) but not to micelles devoid of PG or CL, or those containing the zwitterionic phospholipid phosphatidylethanolamine (PE) **(Fig. 3A, Fig. 4A and Fig. S11)**. Significant CSPs were noted for A74, G120-I128, K131-R133, Q135-L137, V142-S145, I173 and S178-V180. The perturbed residues were mapped to the structure, revealing a single extensive binding site centered on BON2:α1 that was sufficiently large to contact several lipid molecules **(Fig. 3A)**. A dissociation constant (K_d_) of ~80 μM was measured, indicating a tighter association of DolP with PG micelles than other lipoproteins, such as the BAM complex component BamE ^22^. No lipid interaction was seen for any BON1 domain residue, emphasizing the specialized role of BON2, which not only differs from DolP BON1, but also from the BON domains of OsmY and Kbp **(Fig. S8A)**. Analysis of the electrostatic surface reveals a large negative surface potential on BON1:α1, which is absent in BON2:α1 and may act to repel BON1 from PG, whilst BON2:α1 uniquely harbors an aromatic residue W127 in the observed PG binding site **(Fig. S10)**.

**Fig. 3.**
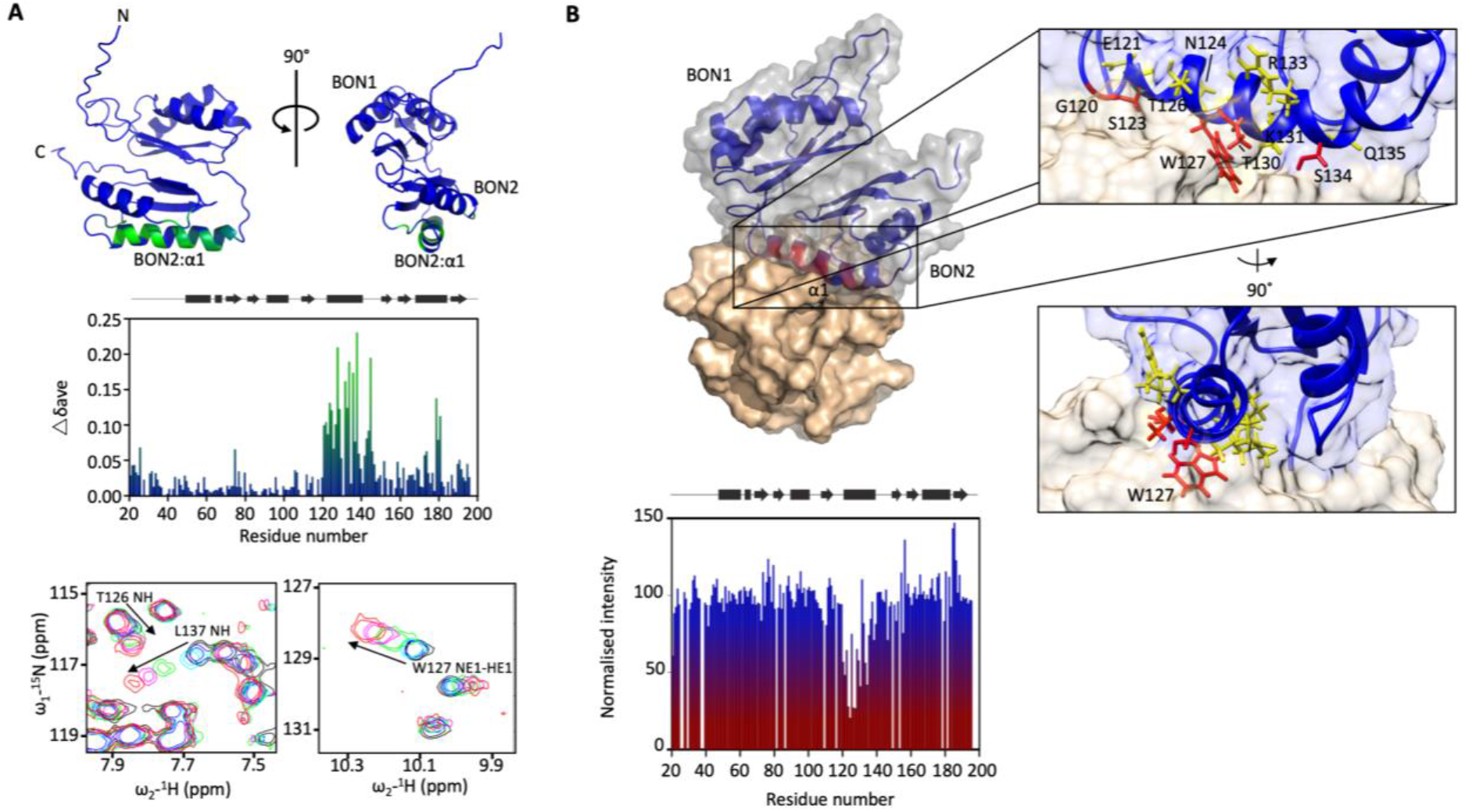
DolP BON2:α1 binds phospholipid. **A.** DolP ribbon structure highlighting residues exhibiting substantial CSPs (Δδ_ave_) upon DHPG micelle interaction. The histogram shows the normalised perturbations induced in each residue’s amide signal when DHPG (40mM) was added to DolP (300 µM). Examples of significant CSPs are shown. **B.** Histogram showing intensity reductions of H_N_ signals of DolP induced by adding 5-doxyl PC and DMPG into DPC/CHAPs micelles and the corresponding structure of a representative DolP-micelle complex calculated using CSPs and doxyl restraints using the program HADDOCK. Only the BON2:α1 helix is observed making contact with the micelle surface. No corresponding interaction of the BON1:α1 helix is observed. Zoom panels show burial of BON2:α1 into the micelle. The side chains of DolP residues that intercalate between the acyl chains (G120, S123, W127, T130 and S134) are coloured red. The side chains of residues that buttress the interface (E121, N124, T126, I128, K131, R133 and Q135) are coloured yellow. DolP is shown in blue and the phospholipid micelle is shown in tan.

**Fig. 4.**
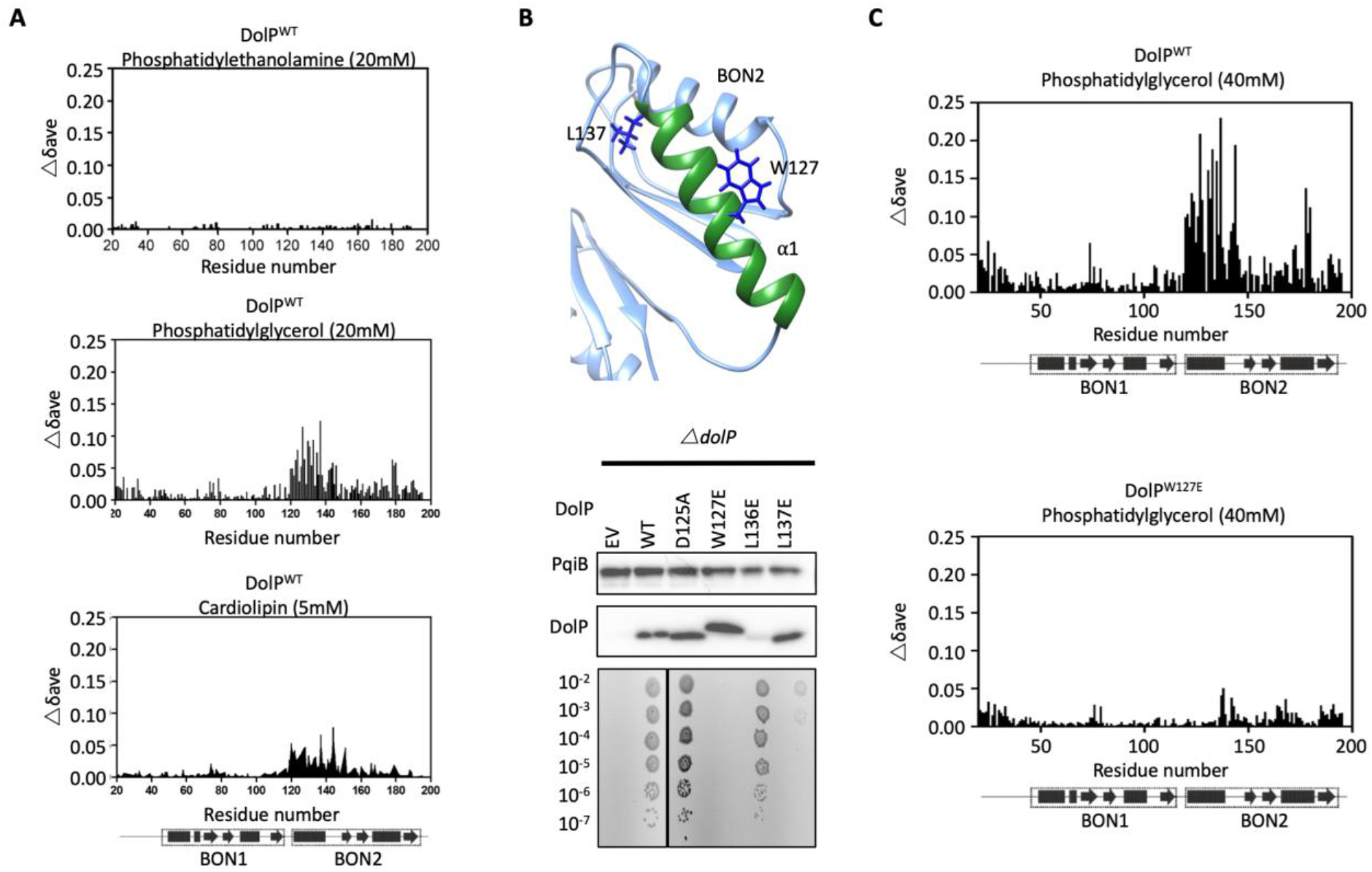
DolP specifically recognizes anionic phospholipid via BON2:α1. **A.** Histograms showing the normalised CSP values observed in ^15^N labelled DolP (300 μM) amide signals in the presence of 20 mM 1,2,-dihexanoyl-sn-glycero-3-phosphethanolamine, 20 mM 1,2-dihexanoyl-sn-glycero-3-phospho-(1’-rac-glycerol) and 5 mM cardiolipin. **B.** Mutagenesis of the BON2:α1 helix residues identified by CSPs. The positions of W127 and L137 are indicated as sticks. Western blots of total protein extracts show plasmid-mediated expression of DolP in *E. coli* Δ*dolP* after site-directed mutation of amino acid residues. The empty vector (EV) control is labelled and WT represents wild type DolP. Colony growth assays of *E. coli* Δ*dolP* complemented with DolP mutants reveal which residues are critical for the maintenance of OM barrier function. The presence of the protein PqiB was used as a control. **C.** Histograms showing the normalised CSP values observed in ^15^N labelled DolP^WT^ or DolPW^127E^ mutant (300 μM) amide signals in the presence of 40 mM 1,2-dihexanoyl-sn-glycero-3-phospho-(1’-rac-glycerol).

As the BON2 domain contained a particularly large PG-specific interaction site, we sought to resolve the micelle-complexed structure of mature DolP. Intermolecular structural restraints were obtained from paramagnetic relaxation enhancements (PRE) obtained by incorporating 5-doxyl spin-labelled phosphatidyl choline (PC) and 1,2-dimyristoyl-*sn-*glycero-3-phospho-(1’-rac-glycerol) (DMPG) into a *n*-dodecylphosphocholine (DPC) micelle and by measuring CSPs. The complexed structure was calculated using HADDOCK^23^ with 18 PRE distance restraints and side chains of the 25 chemical shift perturbations, with final refinement in water **(Fig. 3B)**. The amino acids G120-T130 and V132-S139 were observed to insert into the micelle interior based on the PRE and CSP data. This reveals an unprecedented burial of the BON2:α1 helix, which spans the entirety of the L119-S139 sequence. The protein-micelle interface buries 1358 ± 316 Å^2^ and to our knowledge represents the most extensive structured surface of a membrane:protein interface resolved to date. The surface forms intimate contacts with at least six proximal phospholipid headgroups through an extensive network of highly populated hydrogen bonds and electrostatic interactions. Whilst the side chains of residues G120, S123, W127, T130 and S134 intercalate between the acyl chains, E121, N124, T126, I128, K131, R133 and Q135 buttress the interface **(Fig. 3B)**. This element was also functionally important based on our transposon screen **(Fig. 2E)**, and was further confirmed as being essential by directed mutagenesis. Mutations within the PG-binding BON2:α1 disrupt the function of DolP, the most critical of which are W127E and L137E; W127 is located in the center of the binding site that penetrates deep into the core of the PG micelle, and L137 is located at the periphery of the helix **(Fig. 3B, Fig. 4B and Fig. S13)**. Not only does mutation of W127 lead to loss of function, but introduction of the W127E mutation was shown to abolish binding of DolP to PG micelles as observed by a loss of CSPs within BON2:α1 **(Fig. 4C)**. Notably, the BON2:α1 structure presents an extended α-helix when compared to BON1:α1 **(Figs. S7 and S8)**. The helical extension in BON2:α1 contains the W127 anionic phospholipid-binding determinant of DolP. This further implicates W127, which is absent in BON1 and OsmY, in specialization of DolP BON2 for phospholipid binding.

### Phospholipid binding guides DolP localization to the cell division site

DolP binds anionic phospholipid, which demonstrates sub-cellular localization to sites of higher membrane curvature including the cell poles and division site ^24–26^. To determine if DolP also shows a preference for such sites, we constructed a plasmid expressing a DolP-mCherry fusion and utilizing fluorescence microscopy we observed DolP localized specifically to the cell division site **(Fig. 5A)**. Considering that DolP is non-functional when targeted to the IM **(Fig. S5)**, we investigated if DolP could still localize to the site of cell division when it was mistargeted to the IM; no septal localization was observed **(Fig. S5)**. Next, we tested whether the phospholipid binding activity is also required for division site localization of DolP. We found that introduction of the W127E mutation, which prevents interaction of DolP with PG/CL micelles, abolished division site localization of DolP **(Fig. 5A)**. Considering that W127E not only abolished PG/CL binding, but also division site localization, we concluded that division site localization of DolP was dependent upon binding of DolP to anionic phospholipids, which have previously been shown to be enriched at the division site^25,26^.

**Fig. 5.**
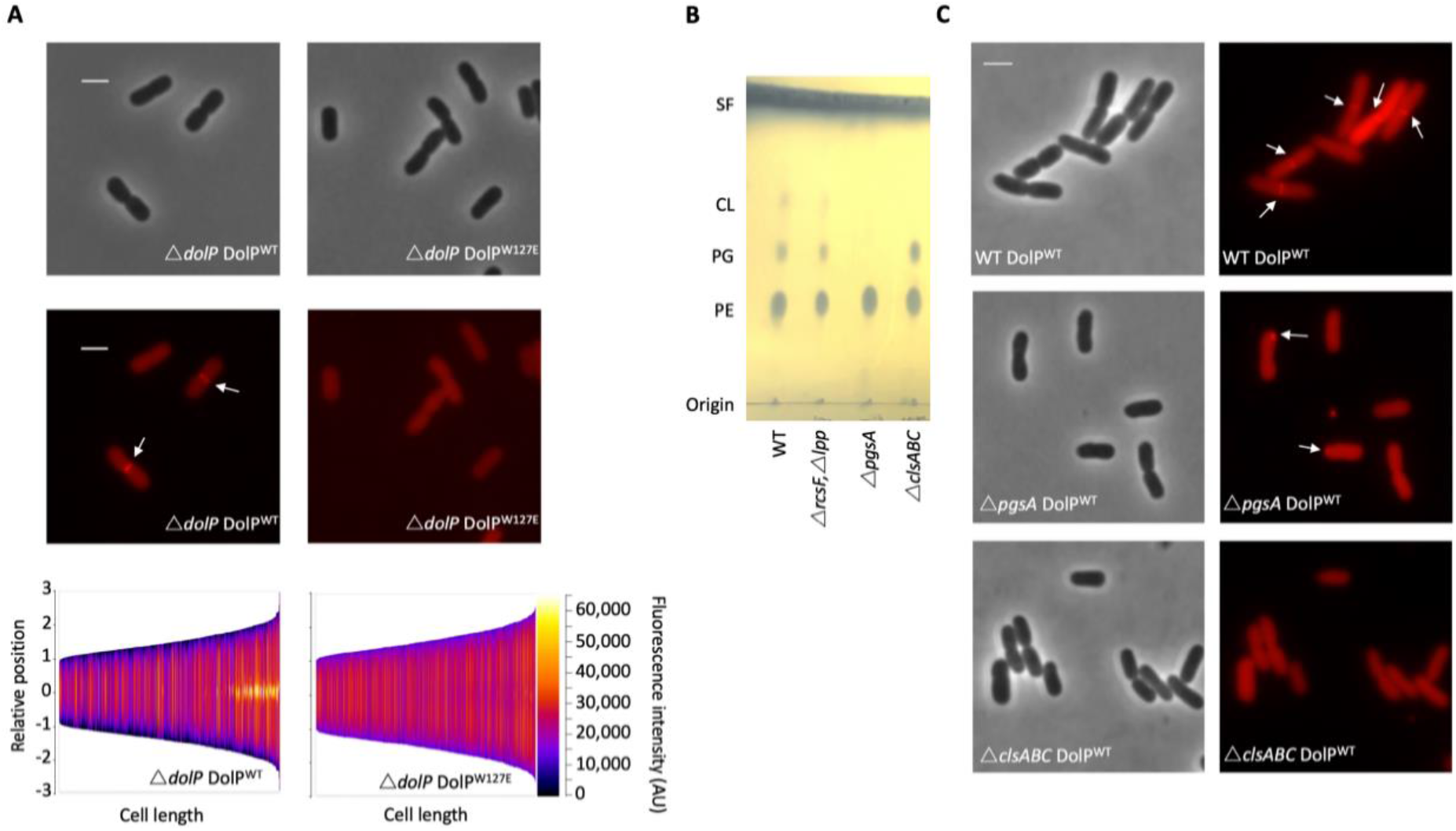
Phospholipid binding is required for DolP recruitment to division sites. **A.** Fluorescence microscopy of Δ*dolP* cells expressing either DolP^WT^::mCherry or DolP^W127E^::mCherry from the pET17b plasmid after growth to mid-exponential phase (OD_600_ ~0.4-0.8). Scale bars represent 2 μM and both phase contrast and the mCherry channel are shown in greyscale and red respectively. White arrows highlight division site localisation of DolP^WT^-mCherry. Demographic representations of the DolP^WT^-mCherry or DolPW^127E^-mCherry fluorescence intensities measure along the medial axis of the cells. Images of >500 cells were analysed using the MicrobeJ software and sorted according to length where the y-axis represents relative cellular position with 0 being mid-cell and 3 or −3 being the cell poles^76^. **B.** Thin layer chromatography of phospholipids extracted from either *E. coli* BW25113 (WT), Δ*rcsF*Δ*lpp*, Δ*rcsF*Δ*lpp*Δ*pgsA* (referred to as Δ*pgsA*) or Δ*clsA*Δ*clsB*Δ*clsC* (referred to as Δcls*ABC*) strains. The *rcsF* and *lpp* genes must be removed in order to prevent toxic build-up of Lpp on the IM in the *pgsA* mutant. Phospholipids were separated using chloroform:methanol:acetic acid (65:25:10) as the mobile phase before staining with phophomolybdic acid and charring. C. Fluorescence microscopy of Δ*pgsA* or Δcls *ABC* cells expressing DolP^WT^mCherry from the pET17b plasmid after growth to mid-exponential phase (OD_600_ ~0.4-0.8). White arrows highlight DolP-mCherry mislocalisation.

To confirm this result we analyzed DolP localization in a strain that lacks all three cardiolipin synthases and is defective for cardiolipin synthesis, which was confirmed by phospholipid extraction and thin layer chromatography **(Fig. 5B)**. We observed that DolP localization is perturbed in the CL-strain, with less dividing cells showing localization of DolP to the septum **(Fig. 5C)**. These effects are further exacerbated in a strain that does not synthesize the major cell anionic phospholipids phosphatidylglycerol or cardiolipin, as confirmed by phospholipid extraction and thin layer chromatography **(Fig. 5B)**. Loss of both phosphatidylglycerol and cardiolipin synthesis worsened the severity of the localization defect with less septal localization and a significant proportion of cells showing mislocalization of DolP to patches at the cell poles **(Fig. 5C)**. Taken together these data demonstrate that DolP localization to the division site is dependent upon interaction with anionic phospholipid *via* BON2:α1, and that this interaction and the sub-cellular localization are required for DolP function.

## Discussion

We have revealed the first structure of a dual-BON domain protein, a protein architecture that is widely conserved among bacteria and therefore provides insight into a diverse range of proteins acting in different organisms. We also report the first evidence for direct binding of lipids by BON domains. We show that DolP BON2 demonstrates specificity for the anionic phospholipids PG and CL, which have previously been shown to localize to sites of higher membrane curvature including the cell poles and division site ^24–26^. Interestingly, we detected no phospholipid binding for DolP BON1, which lacks the key W127 phospholipid interaction residue. This key residue is also lacking in the other periplasmic BON domain-containing protein in *E. coli*, OsmY. Thus, we have demonstrated a specialized role for DolP in the cell and our data suggests BON domains are not generalist phospholipid binding domains, as was suggested previously^18^.

Here we show for the first time that localization of DolP to the cell division site is dependent upon recognition of anionic phospholipids by DolP BON2. To our knowledge, this is the only example of this mechanism of localization to the bacterial division site ^27^. Considering anionic phospholipids also accumulate at the old pole, the question of how DolP specifically recognizes the division site remains. We hypothesize that DolP prefers the site of higher positive (convex) curvature found only at the inner leaflet of the OM cell division site *in vivo* and in the PG micelles used in this study. Previous evidence has shown that inhibition of cell constriction, by the addition of cephalexin, also prevents DolP localization to future division sites^12^. This indicates that DolP may require cell constriction for localization to the division site, therefore lending support to the hypothesis that DolP may recognize membrane curvature. An alternative explanation is that the phospholipid binding mode of DolP may trigger interaction with some as yet unidentified division site localized protein partner, but no obvious candidates are offered by published envelope interactome data^13,14^. Nevertheless, these data reveal that DolP function is dependent on localization to the division site through phospholipid binding and localization to the OM through its N-terminal lipid anchor. The model of DolP localization to the cell division site proposed here also provides some evidence that anionic phospholipids localize to sites of high membrane curvature in the OM. While this has been shown for whole cells^24,26^, and the IM through the use of spheroplasts^25^, to our knowledge, no such observation has yet been made for the OM directly. Considering that the OM is significantly different from the IM and is depleted of PG and CL by comparison^28^ **(Fig S10B)**, the localization of these lipids to sites of negative curvature could be further enhanced by the relative scarcity of these lipids in the OM and this warrants further study.

We have not found a direct mechanism through which DolP maintains OM integrity. Previous protein:protein interaction studies captured DolP as a near neighbour of two components of the Bam complex, BamD and BamE ^13,14^. Consistent with this, *dolP* shows synthetic lethality with the gene encoding the periplasmic chaperone SurA, leading to suggestions of a role for DolP in OMP biogenesis ^29–31^. However, we were unable to demonstrate a direct interaction between DolP and the BAM complex, and no such interaction has been seen in the extensive studies evaluating the subunit composition and multimeric states of the BAM complex^32–35^ or in similar studies in *N. meningitidis*^11^. However, while this is in agreement with the fact that DolP is localized to the division site, whereas the Bam complex is uniformly present across the cell surface^34^, it does not rule out potential transient interactions. Previous observations revealed that the OM is a rigid structure ^36^ that this membrane rigidity stabilizes assembly precincts ^34^, and that the activity of the BAM complex is sensitive to increases in membrane fluidity^21^. We suggest that the increased membrane fluidity of the *dolP* cells, demonstrated here, provides a challenging environment for assembly precincts to be maintained. We hypothesize that DolP, perhaps through interactions with peptidoglycan amidases^12^, might also modulate peptidoglycan remodeling in such a way as to minimize the clash between the periplasmic components of the assembly precinct and the cell wall, which might be exacerbated in regions of high membrane curvature.

In conclusion, this study reports for the first time the direct binding of lipid by BON domains and a new mechanism of protein division site localization. The indirect link between DolP and the general machinery responsible for outer membrane biogenesis adds to the recently described role of DolP in the regulation of cell wall amidases during division, therefore potentially placing DolP at the interface between envelope biogenesis processes^12^. The demonstration that loss of DolP increases sensitivity to antibiotics and membrane disrupting agents, in addition to the decrease in virulence *in vivo*, and an increase of the efficacy of the *N. meningitidis* vaccine, suggests DolP will provide a useful starting platform for antimicrobial design based on the disruption to regulation of multiple envelope biogenesis mechanisms ^10,37,38^.

## Acknowledgements

This work was supported by the BBSRC (I.R.H. and M.O.: BB/M00810X/1 and BB/L00335X/1; T.J.K. BB/P009840/1), NSERC RGPIN-2018-04994 and Campus Alberta Innovation Program (RCP-12-002C) (M.O.). We would like to thank Georgia L. Isom and Catherine A. Wardius for technical assistance in the laboratory. We thank Professor Corinne Spickett for use of mass spectrometry facilities for phospholipid analyses. We also thank Professor Jeff Cole for critical advice in development of the project.

## Methods

### Bioinformatic analyses

The BON domain profile was obtained from Pfam http://pfam.sanger.ac.uk/^39^ and used as input for HMMER (hmmsearch version 3.1)^40^ against the Uniprot database (http://www.uniprot.org, release 06032013) with an inclusion cutoff of E = 1 without heuristic filters. Sequence redundancy for clustering analysis was minimised using the UniRef100 resource of representative sequences; clustering was performed with the mclblastline program^41,42^ based on the e-value obtained by a BlastP run of all-against-all. Optimal settings for the mcl clustering were manually determined, clustering was performed at an e-value cutoff of 1E-2 and an inflation parameter of 1.2 using the scheme ^7^ setting implemented in mcl. The resulting clusters were matched back to the proteins originally recovered by the HMMER search, and the number of proteins, as well as the number of matched organisms, are summarised for each phylum or subphylum in **Table S1**. UniProt accession numbers of all proteins according to their clusters are given in **Table S2**. The domain annotation was obtained from the InterPro database^42^. For cluster representation **(Fig. 1)**, the program CLANS^43^ was used under the default settings. Clusterings with CLANS was based on a subset of OsmY-, DolP- and Kbp-like proteins identified as described above; the respective accession numbers are given in **Table S5**. Pairwise alignment similarity values were analysed at the Protein Information Resource site (PIR; http://pir.georgetown.edu/).

### Plasmids, bacterial strains and culture conditions

*E. coli* BW25113 was the parental strain used for most investigations. *E. coli dolP::kan*, *osmY::kan* and *kbp::kan* mutants were obtained from the KEIO library^44^ and the mutations transduced into a clean parental strain. *E. coli Δ dolP* was created by resolving the Kan^R^ cassette, as previously described^45^. *E. coli* BW25113 Δ*pgsA* was constructed first by transfer of the *rcsF::aph* allele from the Keio library into *E. col* BW25113 and removal of the *kanR* cassette. The *lpp:aph* allele was then introduced into the Δ*rcsF* strain and the cassette removed by the λ-Red recombination method of Datsenko and Wanner, due to the presence of Lpp being toxic in the absence of phosphatidylglycerol^45–47^. Finally, the same method was utilized to create the Δ*pgsA* strain (Δ*rcsF*,Δ*lpp*,Δ*pgsA*) The genes encoding DolP and OsmY were amplified from *E. coli* BW25113 and cloned into pET17b to create pDolP and pOsmY. Orthologous sequences from *S. enterica, V. cholera, N. meningitidis, H. influenza* and *P. multocida* were synthesized and cloned into pET17b to create the plasmids pSe, pVc, pNm, pHi and pPm, respectively. To create pDolP^pelB^, the gene encoding DolP was synthesized but with nucleotides encoding the PelB signal sequence in place of the native signal sequence and without Cys19 to relieve the possibility of acylation; this plasmid was constructed in pET26b+ such that the protein had a C-terminal His-tag. In addition, to create p(OM)OsmY the gene encoding OsmY was synthesized but with nucleotides encoding the native DolP signal sequence and Cys19 N-terminal acylation site in place of the native OsmY signal sequence. The latter plasmid was constructed in pET17b. The pET17b-*dolP::mCherry* plasmid was constructed to contain an 11 amino acid flexible linker and a codon optimized mCherry gene at the 3’ end of the *dolP* gene. Gene synthesis was performed by Genscript®. The pet20b+-*wbbL* plasmid for restoring O-antigen synthesis in *E. coli* K-12 was previously described^48^. Single point mutations were generated by using Quickchange II according to manufacturer’s instructions. All constructs were confirmed by DNA sequencing. Strains were routinely cultured on LB agar and LB broth. Linker scanning mutagenesis was performed with an Ez-Tn5 kit (Epicentre®) as previously described^49^.

### Analysis of membrane lipid content

Cell envelopes of *E. coli* were separated by defined sucrose density gradient separation, precisely as described previously following cell disruption by 3 passes of the C3 emulsiflex (Avestin)^50,51^. Samples were generated from 2 L of cells grown to an OD_600_ 0.6-0.8, with the final volumes for washed membranes being 1 ml, which were stored at −80°C until analysis. Lipids were extracted by the Bligh-Dyer method^52^ from purified membranes as described previously^50^. Methanol and chloroform were added to the samples to extract the metabolites using a modified Bligh-Dyer procedure^53^ with a final methanol/chloroform/water ratio of 2:2:1.8. The non-polar layer was extracted and dried under nitrogen before being stored at −80°C until analysis. Samples were re-dissolved in 200 μl chloroform before being separated by thin layer chromatography on silica gel 60 plates with the mobile phase as chloroform:methanol:water at the following ratio: 65:25:10. Lipids were visualized by staining with phosphomolybdic acid. Analysis of lipid samples by mass spectrometry was completed as described previously ^54^. The differences were as follows: lipid extracts were diluted 10x or 20x into starting LC solvent the LC-MS/MS run directly. Normalization was completed by taking the ion intensity of each phospholipid relative to the total ion count.

### Biochemical analyses

Cellular fractions were prepared as described previously^55^. Cellular fractions and purified proteins were electrophoresed on 12 or 15% SDS-PAGE gels and stained with Coomassie blue or transferred to a polyvinylidene difluoride (PVDF) membrane for Western immunoblotting as previously described^56^. Loading consistency was confirmed by immuno-blotting with anti-BamB or anti-PqiB antiserum where possible. Protease shaving assays were described previously^57^. Proteins were localized by immunofluorescence as described previously^56^. Analytical ultracentrifugation was performed as described previously^22^. For proteomic analysis of OM protein content, OM fractions purified by defined sucrose gradient centrifugation were digested with trypsin using the FASP method^58^. Primary amines in the peptides were then dimethylated using hydrogenated or deuterated formaldehyde and sodium cyanoborohydride. Labelled peptides were mixed, separated into 15 fractions by mixed-mode reverse-phase/anion exchange chromatography, the fractions lyophilized and each analysed with a 90 minute LC-MS/MS run using a Bruker Impact Q-TOF mass spectrometer. Data was searched against forward and randomized *E. coli* sequence databases using MASCOT and filtered at 1% FDR. Quantitation was based on the extracted ion chromatograms of light/heavy peptide pairs. DolP was investigated for binding partners using immunoprecipitation assays as described previously. Briefly, *E. coli ΔdolP*, and isogenic strains containing pDolP^pelB^ or plasmid containing a His-Tagged version of BamA were grown in LB media to an OD_600_ of ~0.6 and harvested by centrifugation. Cells were resuspended in PBS with Triton X-100 supplemented with lysozyme and Benzonase nuclease. Cells were lysed and clarified by centrifugation. The lysate was incubated with Ni-NTA agarose (Qiagen) or appropriate antibodies. Precipitated proteins were analysed by Western immunoblotting.

### NMR spectroscopy

Experiments were carried out at 298 K on a Varian Inova 800 MHz spectrometer equipped with a triple-resonance cryogenic probe and *z*-axis pulse-field gradients. Isotope labelled DolP (^15^N ^13^C) with its N-terminal cysteine replaced was used at a concentration of 1.5 mM in 50 mM sodium phosphate (pH 6), 50 mM NaCl and 0.02% NaN3 in 90% H_2_O/10% D_2_O. Spin system and sequential assignments were made from CBCA(CO)NH, HNCACB, HNCA, HN(CO)CA, HNCO, HN(CA)CO, H(C)CH TOCSY and (H)CCH TOCSY experiments^59^. Spectra were processed with NMRPipe^60^ and analyzed with SPARKY^61^.

### Structure calculations

Interproton distance restraints were obtained from ^15^N- and ^13^C-edited NOESY-HSQC spectra (T_mix_=100 ms). PRE restraints were obtained by adding 10 mM DPC/3.33 mM CHAPS micelles spiked with 1 mM DMPG and 0.185 mM 5-doxyl 1-palmitoyl-2-steroyl-sn-glycero-phosphocholine (Avanti, Polar Lipids, Alabaster, AL, USA) to ^15^N-labelled DolP (300µM) and by standardising amide resonance intensities to those induced by spiking instead with unlabelled dipalmitoyl phosphocholine (Avanti Polar Lipids). Backbone dihedral angle restraints (ϕ and ψ) were obtained using TALOS from the backbone chemical shifts^62^. Slowly exchanging amides were deduced from the ^1^H ^15^N SOFAST-HSQC^63^ spectra of protein dissolved in 99.96% D_2_O. The structure was calculated iteratively using CANDID/CYANA, with automated NOE cross-peak assignment and torsion angle dynamics implemented^64^. A total of 20 conformers with the lowest CYANA target function were produced that satisfied all measured restraints. Aria1.2 was used to perform the final water minimization^65^. Structures were analysed using PROCHECK-NMR^66^ and MOLMOL^67^. Structural statistics are summarized in **Table S3**.

### Lipid interactions

Ligand binding to 300 μM ^15^N-DolP in 50 mM sodium phosphate (pH 6), 50 mM NaCl and 0.02% NaN_3_ in 90% H_2_O/10% D_2_O was monitored by ^1^H^15^N-HSQCs at concentrations of 0–40 mM of either DHPG or DHPE (c.m.c., ~7 mM). The DPC-DMPG: DolP complex was calculated by HADDOCK^23,68^. A total of 18 paramagnetic relaxation enhancements restrained the distances between the micelle centre and the respective NH groups to 0-20 Å, with CSPs defining the flexible zone. The top 200 models were ranked according to their experimental energies and statistics derived from the 20 lowest energy conformers were reported **(Table S6)**.

### Small angle X-ray scattering

Synchrotron SAXS data of DolP were collected at the EMBL X33 beamline (DESY, Hamburg) using a robotic sample changer. DolP concentrations between 1-10 mg/ml were run in 50 mM sodium phosphate (pH 6), 50 mM NaCl and 0.02% NaN_3_. Data were recorded on a PILATUS 1M pixel detector (DECTRIS, Baden, Switzerland) at a sample-detector distance of 2.7 m and a wavelength of 1.5 Å, covering a range of momentum transfer of 0.012 < s < 0.6 Å^−1^ (s = 4πsin(θ)/γ, where 2θ is the scattering angle) and processed by PRIMUS^69^. The forward scattering I(0) and the radius of gyration (R_g_) were calculated using the Guinier approximation^70^ **(Fig. S14)**. The pair-distance distribution function pR, from which the maximum particle dimension (D_max_) is estimated, was computed using GNOM^71^ **(Fig. S14)**. Low resolution shape analysis of the solute was performed using DAMMIF^72^. Several independent simulated annealing runs were performed and the results were analysed using DAMAVER^73^. Back comparison of the DolP solution structure with the SAXS data was performed using the ensemble optimisation method^74^ accounting for flexibility between residues 20-46, 112-118 and 189-195. All programs used for analysis of the SAXS data belong to the ATSAS package^75^.

### Accession codes

Coordinates and NMR assignments have been deposited with accession codes 2mk8 (PDB) and 19760 (BMRB), respectively.

### Cell imaging

Cultures were grown at 37°C to OD_600_ 0.4-0.5. Cells were harvested by centrifugation at 7000 × g for 1 min before being applied to agarose pads, which were prepared with 1.5 % agarose in PBS and set in Gene Frames (Thermo Scientific). Cells were immediately imaged using a Zeiss AxioObserver equipped with a Plan-Apochromat 100x/Oil Ph3 objective and illumination from HXP 120V for phase contrast images. Fluorescence images were captured using the Zeiss filter set 45, with excitation at 560/40 nm and emission recorded with a bandpass filter at 630/75 nm. For localisation analysis and generation of demographs, the MicrobeJ plugin for Fiji was used and >500 cells were used as input for analysis^76^.

### Membrane fluidity assay

Membrane fluidity was measured by use of the membrane fluidity assay kit (Abcam: ab189819) as was described previously except with minor modifications^21^. Specific bacterial strains were grown to stationary phase overnight (~16 hrs) after which cells were harvested by centrifugation, washed with PBS three times and finally labelled with labelling mix (10 μM pyrenedecanoic acid and 0.08% pluronic F-127 in PBS) for 20 minutes in the dark at 25°C with shaking. Cells were washed twice with PBS before fluorescence was recorded with excitation at 350 nm and emission at either 400 nm or 470 nm to detect emission of the monomer or excimer respectively. Unlabelled cells were used as a control to confirm labelling and the *E. coli* BW25113 Δ*waaD* strain was used as a positive control for increased membrane fluidity. Following subtraction of fluorescence from the blanks, averages from triplicate experiments were used to calculate the ratio of excimer to monomer fluorescence. These ratios were then expressed as relative to the parent *E. coli* BW25113 strain.

### Genetic interaction analysis

Genetic interaction assay was performed as described in^77^. For each probed strain, a single source plate was generated and transferred to the genetic interaction plate using a pinning robot (Biomatrix 6). On each genetic interaction assay plate, the parental strain, the single deletion A, the single deletion B and the double deletion AB were arrayed, each in 96 copies per plate. Genetic interaction plates were incubated at 37°C for 12 h and imaged under controlled lighting conditions (spImager S&P Robotics) using an 18-megapixel Canon Rebel T3i (Canon). Colony integral opacity as fitness readout was quantified using the image analysis software Iris^78^. Fitness ratios were calculated for all mutants by dividing their fitness values by the respective WT fitness value. The product of single mutant fitness ratios (expected) was compared to the double mutant fitness ratio (observed) across replicates. The probability that the two means (expected and observed) are equal across replicates is obtained by a Student’s two-sample *t*-test.

### Lipid A Palmitoylation assay

Labelling of LPS, Lipid A purification, TLC analysis and quantification were done exactly as described previously^79^. The positive control was exposed to 25 mM EDTA for 10 min prior to harvest of cells by centrifugation in order to induce PagP mediated palmitoylation of Lipid A^79^. Experiments were completed in triplicate and the data generated was analysed as described previously.

## Supplementary material

**Fig. S1.**
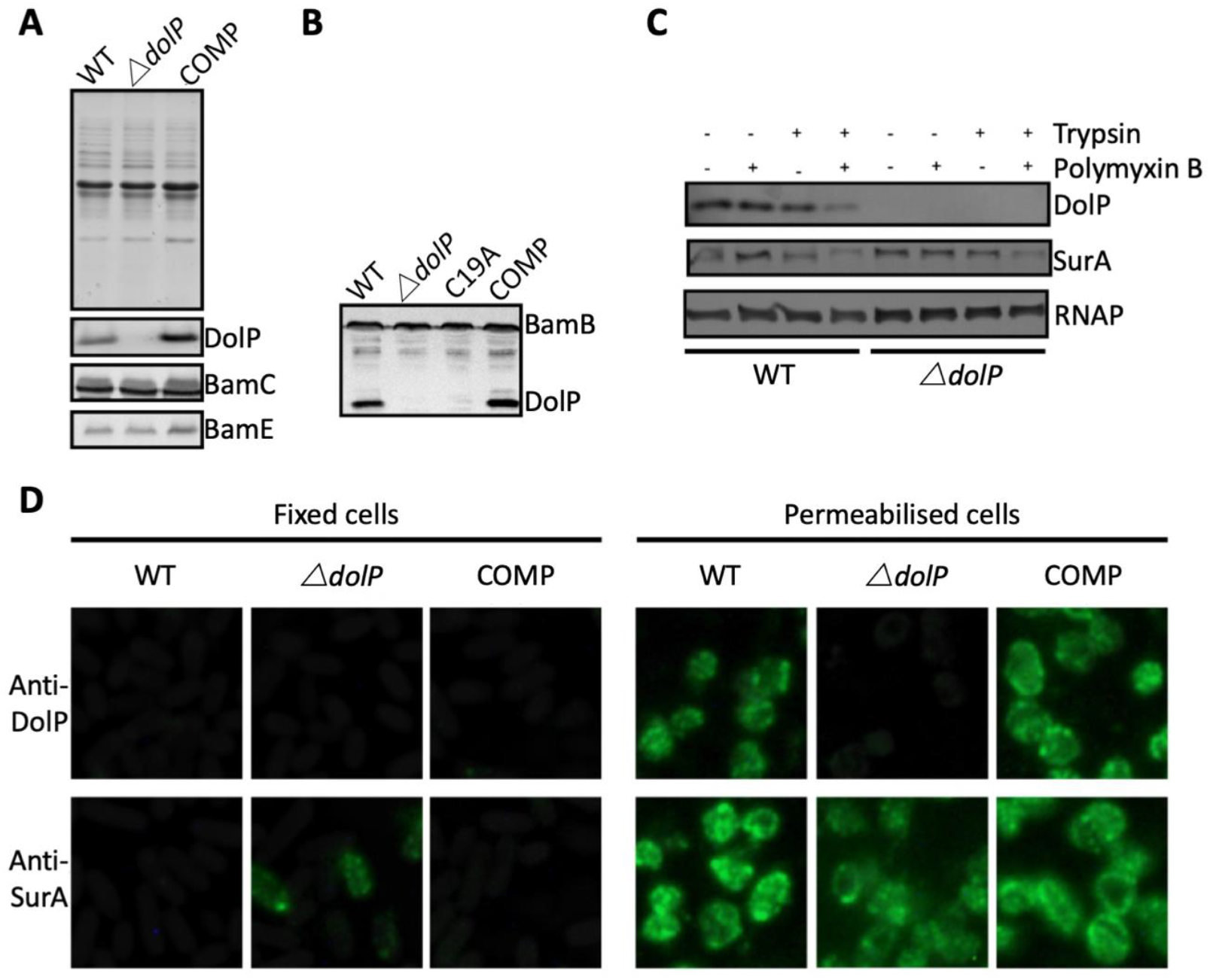
DolP is an OM lipoprotein. **A.** OM fractions of *E. coli* BW25113, an isogenic Δ*dolP* mutant and the complemented mutant were analysed by SDS-PAGE and Western immunoblotting with antibodies to DolP and the known OM lipoproteins BamC and BamE. DolP is not detected in the mutant but like BamC and BamE is found with the membrane fraction. **B.** Western immunoblotting of OM fractions from *E. coli* Δ*dolP* complemented with a plasmid (pDolP-C19A) encoding DolP with a point mutation at position C19. **C.***E. coli* cells treated with protease in the presence (+) or absence (-) of polymyxin B. Antibodies to the cytoplasmic RNA polymerase (RNAP) and the periplasmic chaperone SurA were used as controls. **D.** Immunofluorescence photomicrographs of *E. coli* BW25113, an isogenic Δ*dolP* mutant and the complemented mutant. Cells were probed with anti-DolP before and after permeabilisation. Anti-SurA was used as a control.

**Fig. S2.**
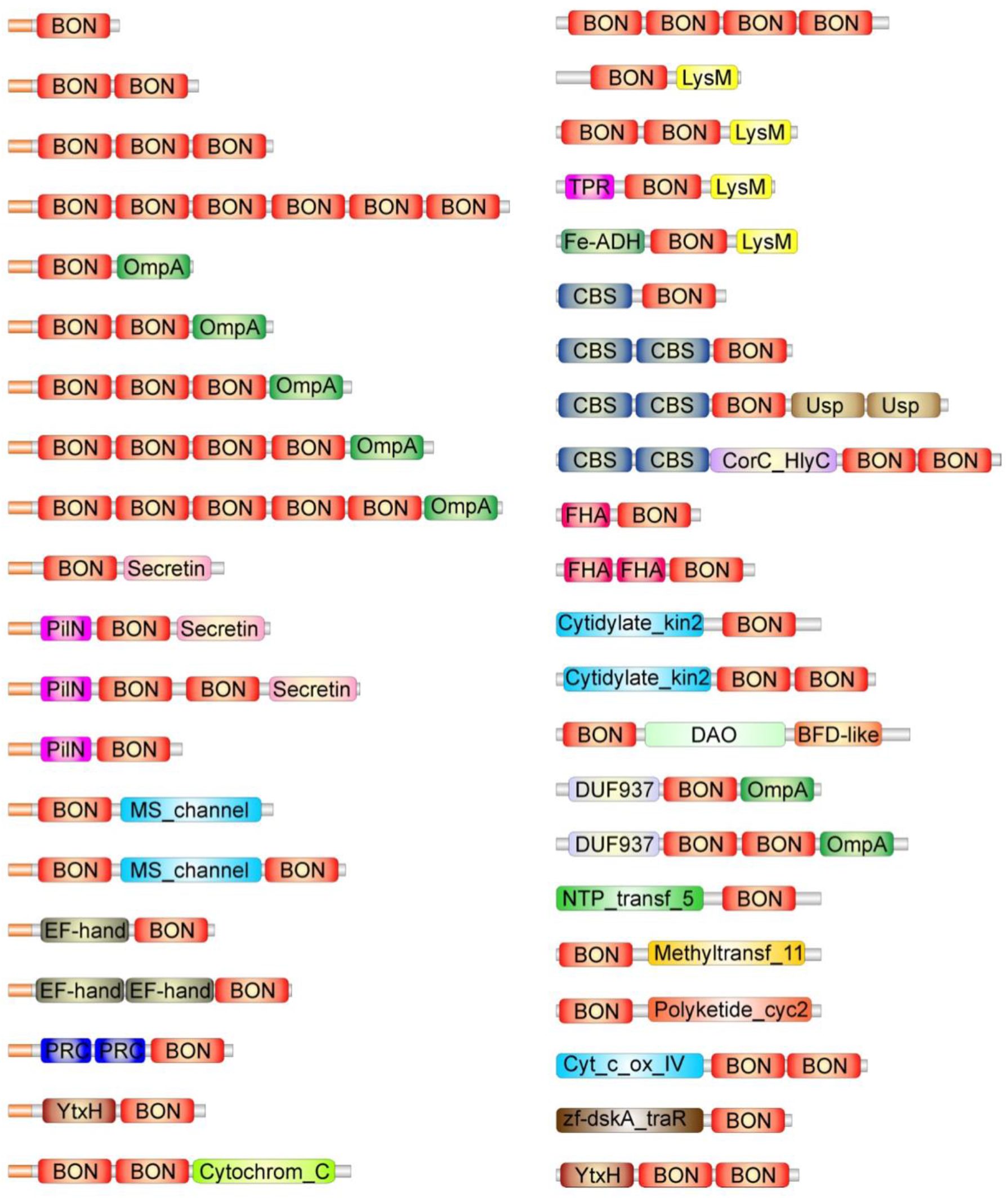
BON domain (Pfam: PF04972) containing proteins. The Pfam database was interrogated for the presence of proteins containing BON domains. BON domains are widely distributed in bacteria and eight major architectures are noted **(Table S1)**. The predominant architecture is that observed for DolP and OsmY where the protein possesses a signal sequence and one or more BON domains. The second major architecture is that observed for Kbp, where proteins possess one or more BON domains and a LysM domain. The other major architectures include associations with Secretin (Pfam: PF00263), CBS (Pfam: PF00571), OmpA (Pfam: PF00691), MS_channel (Pfam: PF00924), FHA (Pfam: PF00498) or cytidylate kinase (Pfam: PF13189) domains. Due to their functions, many of these domains would place their associated BON domains in proximity to cell membranes.

**Fig. S3.**
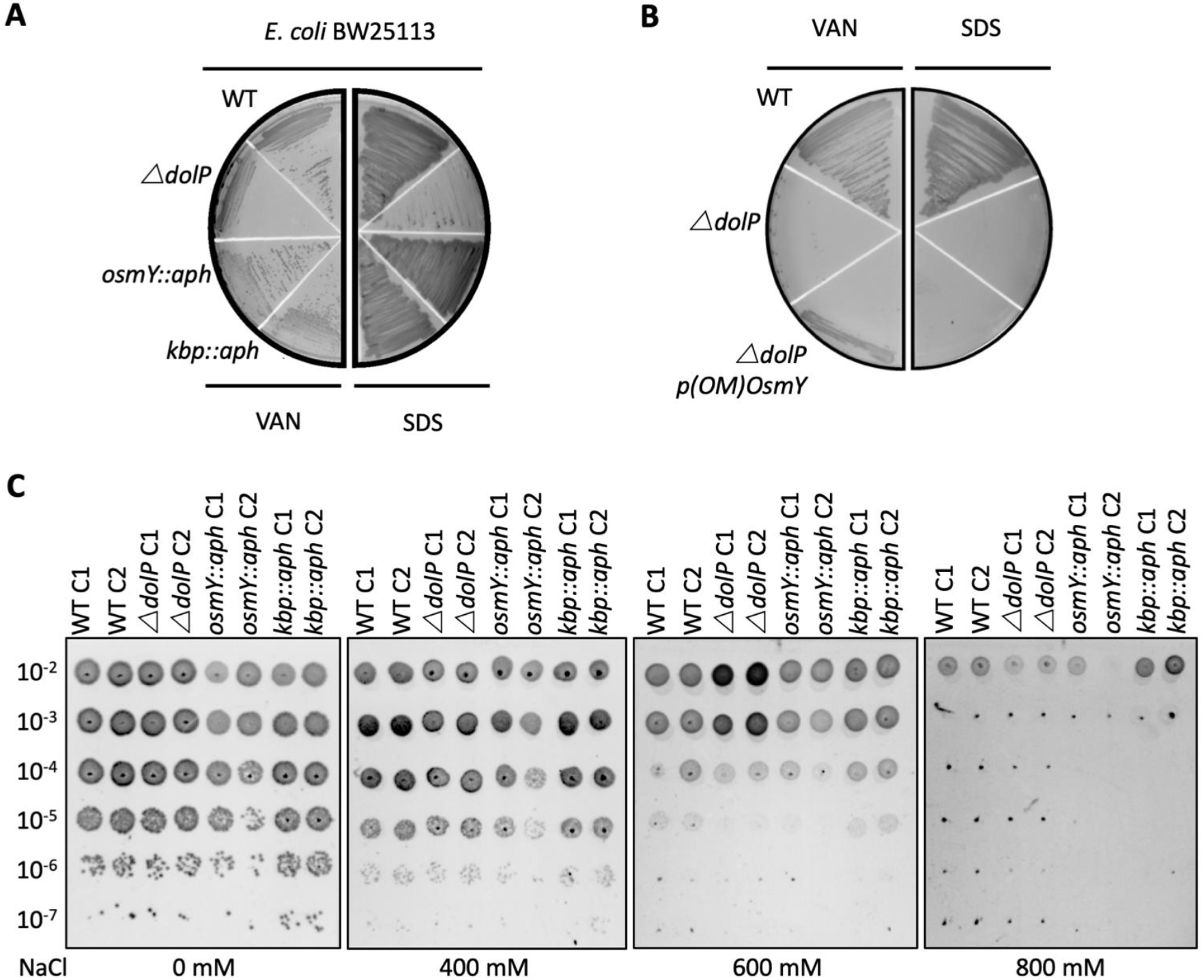
DolP has a distinct function from OsmY and Kbp. The precise functions of Kbp and OsmY are unknown, though both are induced during adaptation to hyperosmolarity^30,80–83^ **A.** Investigation of *osmY* and *kbp* null mutants of *E. coli* revealed neither was sensitive to vancomycin or SDS. Growth phenotypes for mutant isolates lacking BON domain proteins, wild-type strains (WT) or complemented mutants (COMP). Strains were grown on LB agar containing vancomycin (100 μg/ml) or sodium dodecyl sulphate (SDS; 4.8%). **B.** A plasmid encoding a DolP-OsmY chimeric protein composed of the lipoprotein targeting sequence of DolP and the BON domains of OsmY failed to complement the OM defect associated with loss of *dolP*. **C.***E. coli* BW25113 Δ*dolP* is not more susceptible to osmotic stress induced by NaCl than the parent strain as assessed by a serial dilution plate assay. Interestingly, our investigations did not reveal a role for either *kbp* or *osmY* in survival of osmotic stress as the *E. coli* BW25113 parent strain and isogenic *osmY::aph* and *kbp::aph* mutants survived equally well.

**Fig. S4.**
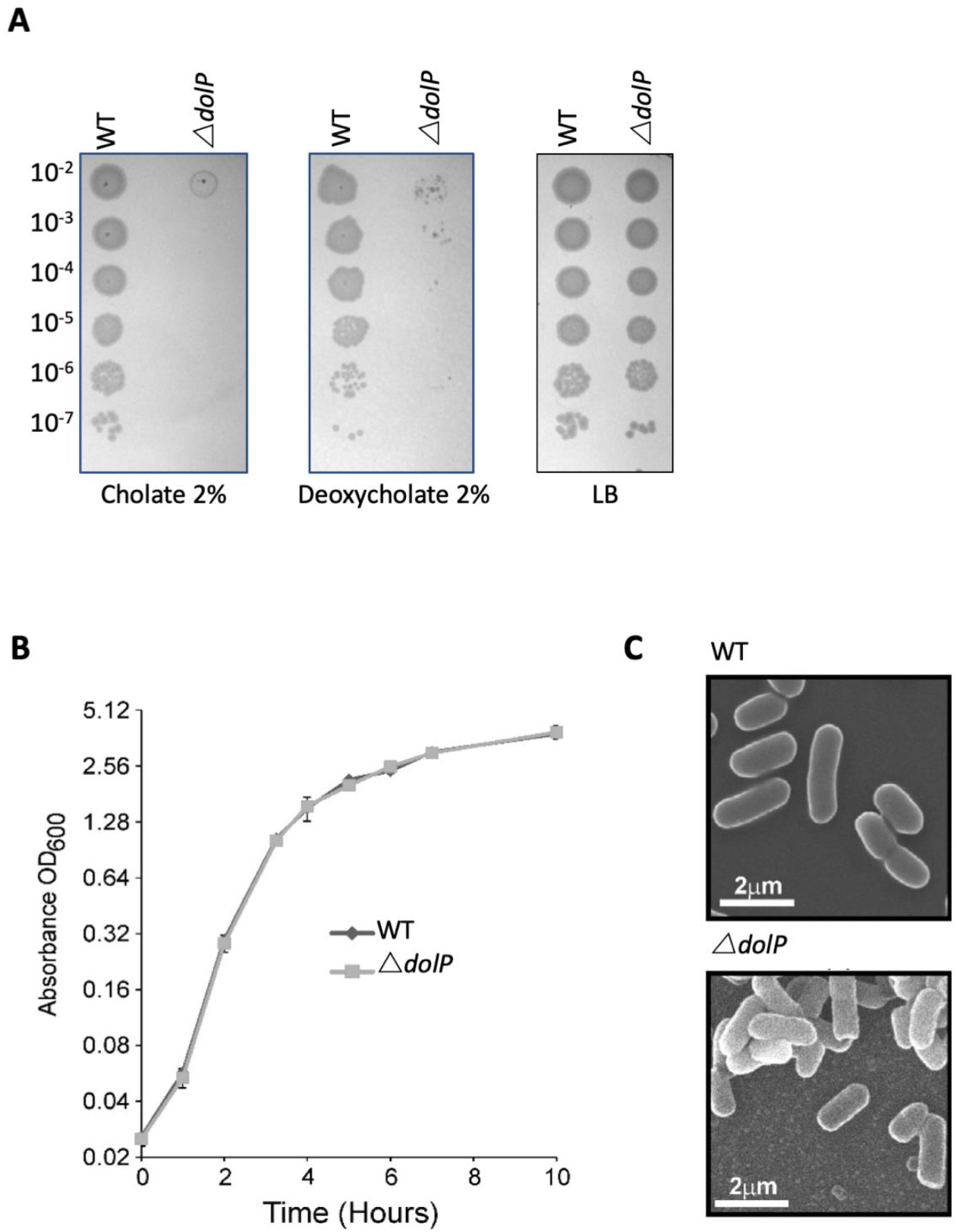
Phenotypes of *E. coli* BW25113 Δ*dolP*. **A.** Mutants lacking *dolP* are sensitive to the anionic detergents cholate and deoxycholate **B.** Mutants lacking *dolP* have growth rates that are indistinguishable from wild-type *E. coli*. **C.** Scanning electron microscopy reveals parental and *E. coli* Δ*dolP* cells have no discernible differences in cellular morphology.

**Fig. S5.**
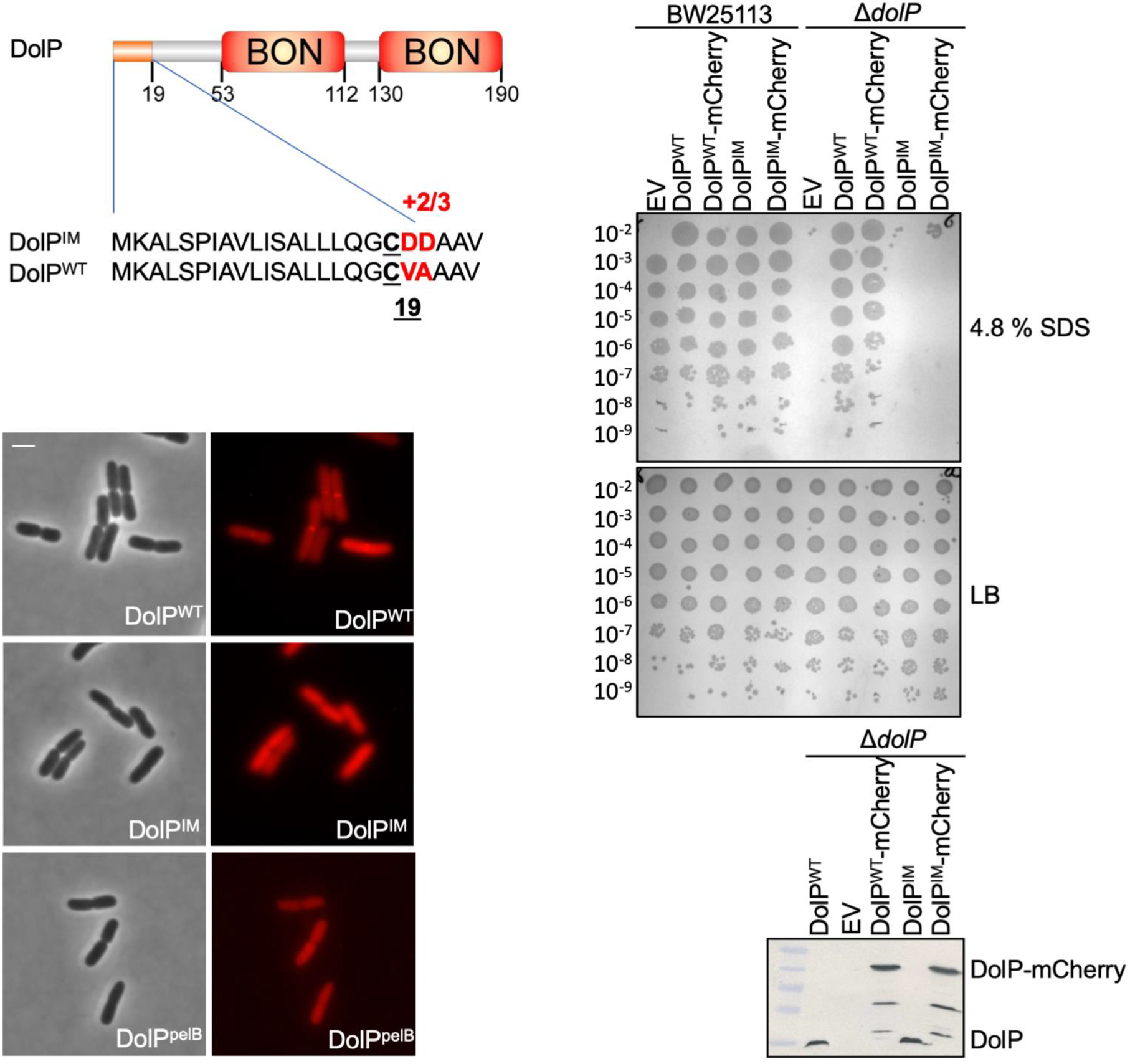
Localisation of DolP to the OM is required for function. The signal sequence and domain architecture of DolP are shown. The sequence changes to pET17b-*DolP^WT^* to create the construct targeting DolP to the IM (pET17b-*dolP*^*IM*^) are shown in red. The signal sequence of *dolP* was also swapped for that of *pelB* in order to create the construct pET17b-*dolP*^*pelB*^ in order to target DolP to the periplasmic space with no modification. Fluorescence microscopy of Δ*dolP* cells expressing either DolP^WT^-mCherry or DolP^IM^-mCherry or DolPpelB-mCherry from the pET17b plasmid after growth to mid-exponential phase (OD_600_ ~0.4-0.8). Scale bars represent 2 μM and both phase contrast and the mCherry channel are shown in greyscale and red respectively. The capacity of DolP^WT^, DolP^IM^, DolP^WT^-mCherry or DolP^IM^-mCherry to complement the Δ*dolP* mutant sensitivity phenotype was screened by dilution assay on 4.8 % SDS. The expression of each construct was checked by Western blotting of total protein extracts with anti-DolP antiserum.

**Fig. S6.**
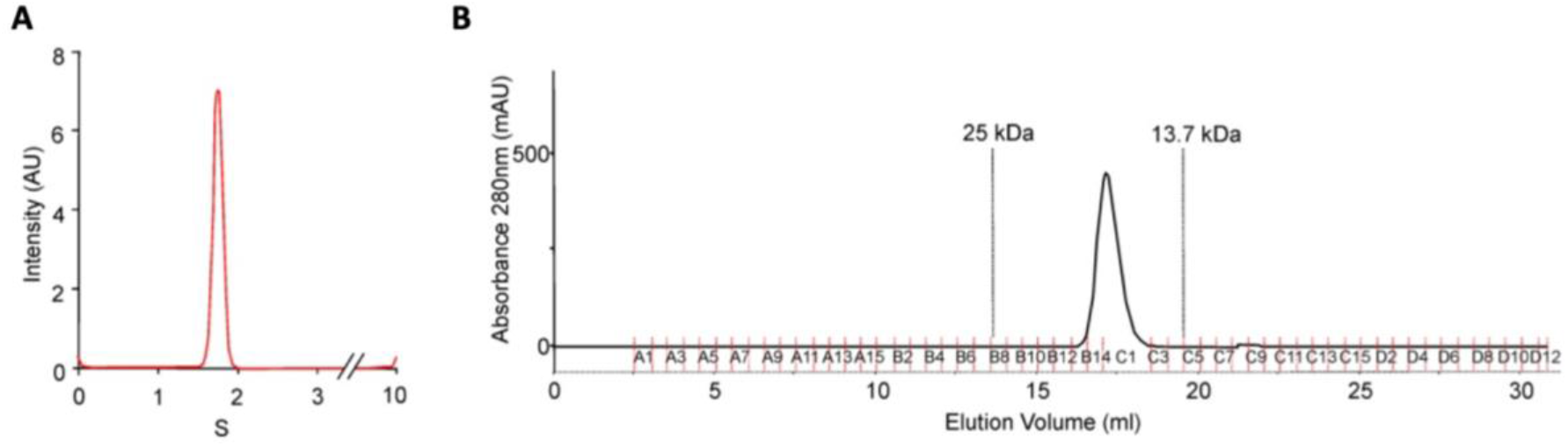
DolP is monomeric **A.** DolP, lacking the site of acylation, was purified and subject to analytical ultracentrifugation. DolP demonstrated a uniform sedimentation velocity consistent with a monomeric species. **B.** Column chromatography of purified DolP revealed that it had an elution profile consistent with a single monomeric species.

**Fig. S7.**
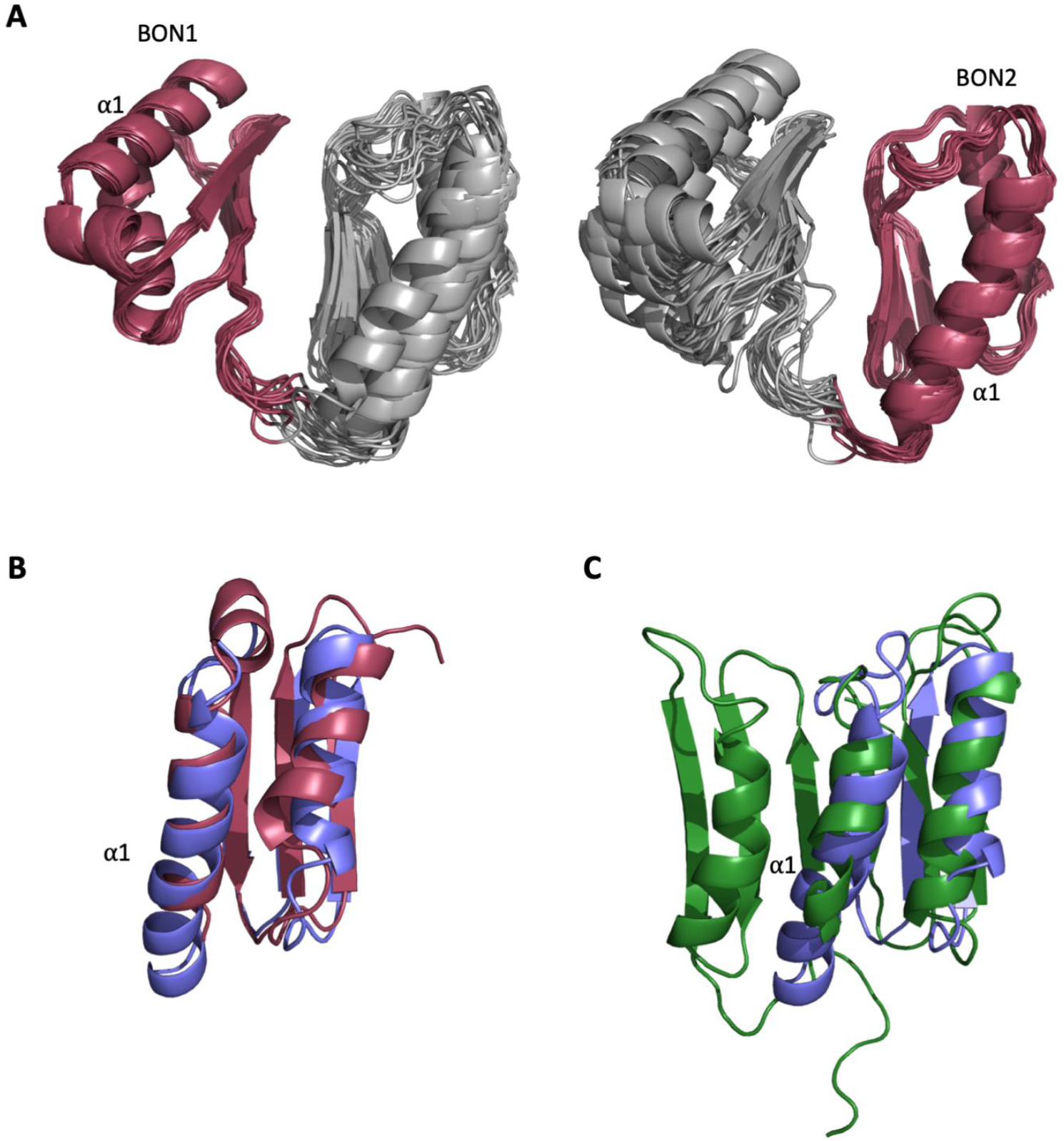
Structural analysis of the DolP BON domains. **A.** The ensemble of the 20 lowest energy structures superimposed to DolP BON1 (N47-I111) and BON2 (G120-T185) domain backbones showing how well the domains superimpose as well as the respective degrees of freedom available to each domain. **B.** Dalilite superposition of DolP BON domains 1 (Red; residues 46-114) and 2 (Blue; residues 117-189). The BON domains are similar except for the double turn extension of the BON2:α1 helix and the presence of the α1’ helix present in BON1 that is absent in BON2. The pairwise RMSD for backbone heavy atoms is 1.8 Å and dalilite Z-score is 8.4. **C.** Superposition of DolP BON2 (Blue) on to the BON subdomain of Rv0899 (OmpATb) (Green; accession code – 2KSM; residues 136-196). For BON2 the pairwise RMSD for backbone heavy atoms was 2.7 Å and the dalilite Z-score was 4.9. Similarly, for BON1 the pairwise RMSD was 2.6 Å and the dalilite Z-score was 5.3.

**Fig. S8.**
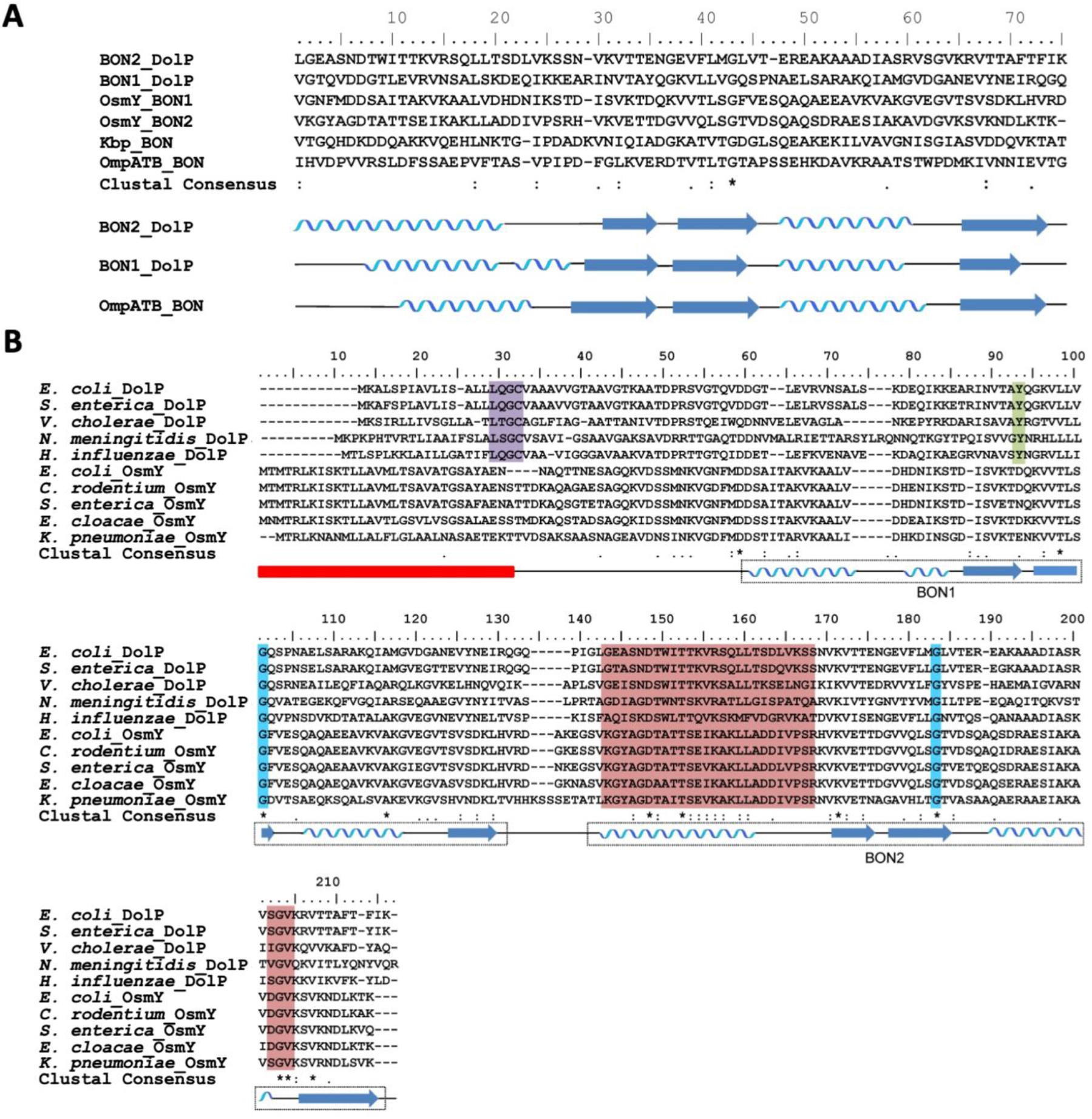
Alignment of DolP sequences from diverse proteobacterial species. **A.** The amino acid sequences of the experimentally derived BON domains of DolP and OmpATb are aligned with the predicted amino acid sequences of the BON domains from Kbp and OsmY. The position of the experimentally derived secondary structure for DolP BON1 and BON2 and OmpATb are depicted below the sequence alignment. **B.** Alignments of the amino acid sequences of DolP and OsmY from various Gram-negative bacteria. The positions of the experimentally-derived secondary structural elements of *E. coli* DolP are depicted below the sequence alignment. The signal sequence is depicted by the red box. The Lipobox associated with recognition by LspA and acylation is highlighted in purple. The conserved glycine residues are highlighted in blue and the tyrosine residue associated with interdomain interactions is highlighted in green. Residues showing CSPs are highlighted in pink.

**Fig. S9.**
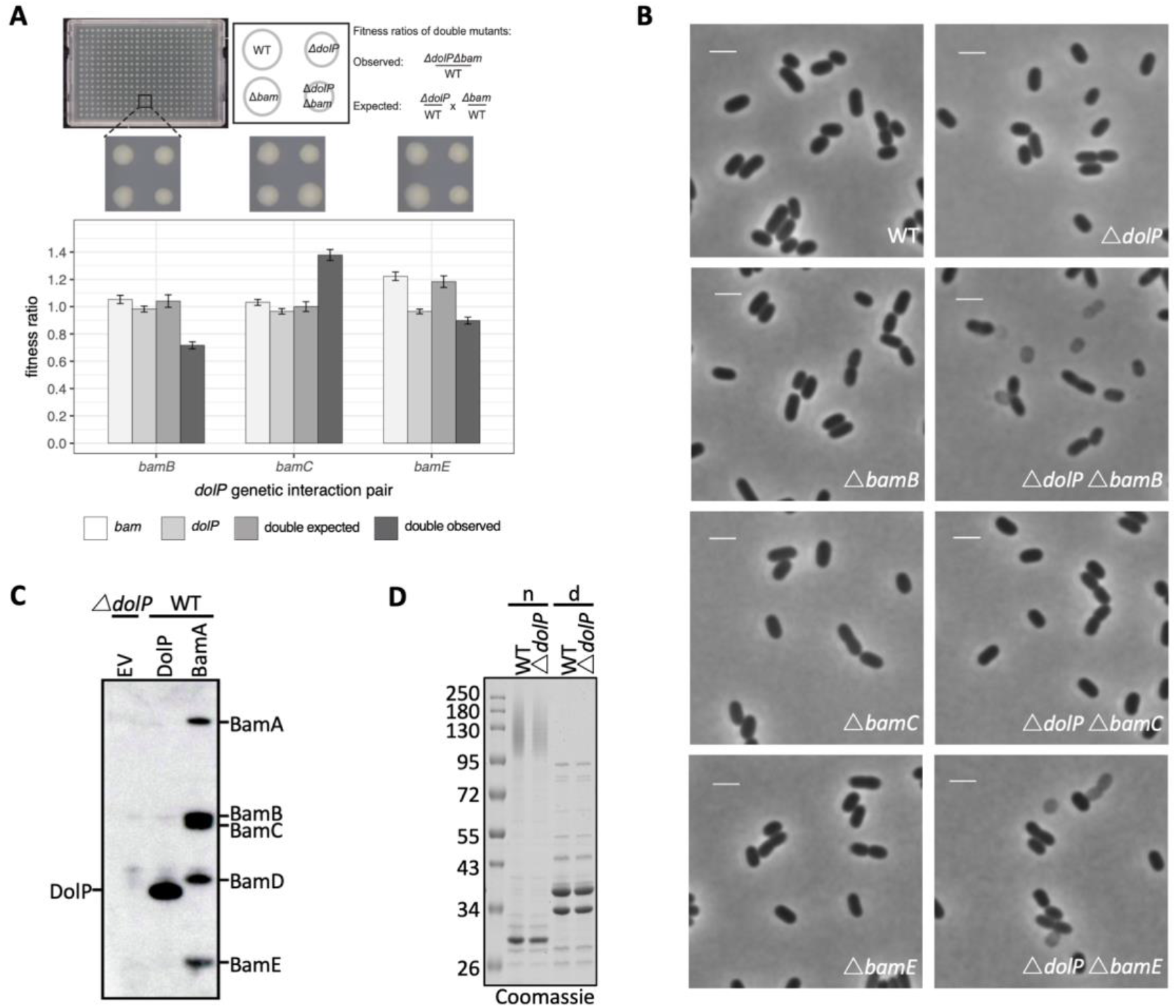
A. *dolP* genetically interacts with the genes encoding the non-essential BAM complex accessory lipoproteins. Strains were arrayed on LB Lennox agar plates using a Biomatrix 6 replicator. Genetic interaction plates were incubated for 12h at 37°C and imaged. An example of a 384-well plate is shown above the graph. Each plate contained a total of 384 colonies consistent of 96 wildtype, single and double mutant clones. Fitness was measured by quantifying colony integral opacity using the image analysis software Iris^78^. Bar plots show the averaged values 96 technical replicates. The error bars represent the 95% confidence interval. **B.** Phase contrast microscopy of WT, Δ*dolP*, Δ*bamB*, Δ*bamC*, Δ*bamE*, Δ*bamB*Δ*dolP*, Δ*bamC*Δ*dolP* and Δ*bamE*Δ*dolP* cells after growth to mid-exponential phase (OD_600_ ~0.4-0.8). Scale bars represent 2 μM. Phase light cells can be observed for the Δ*bamB*Δ*dolP* and Δ*bamE*Δ*dolP* cells. **C.** DolP immunoprecipitation. Whole cell triton X-100 solubilised lysates of *E. coli* BW25113 pDolP^pelB^, pBamA-His and *ΔdolP*, were purified by Ni-NTA affinity chromatography then detected by western blot using anti-DolP and BamA-E antibodies. **D.** Purified OM samples from *E. coli* BW25113 parent (WT) or Δ*dolP* cells were separated by SDS-PAGE, with (d) and without (n) boiling before being visualized by staining with coomassie.

**Fig. S10.**
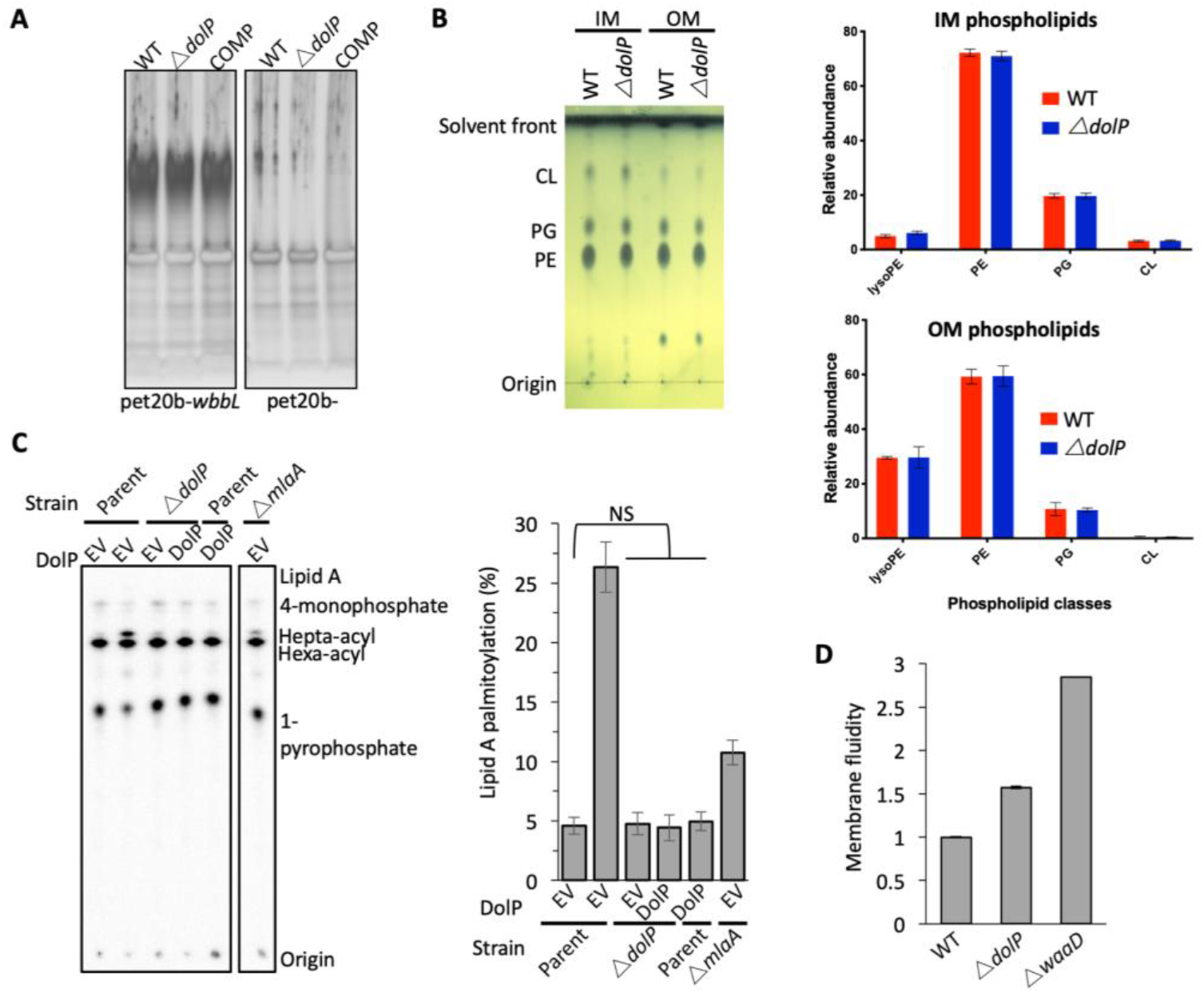
Loss of DolP does not affect membrane lipid profiles. **A.** SDS-PAGE gel showing separation of LPS preparations from *E. coli* BW25113 and *E. coli* BW25113 harboring pET20b-*wbbL* which restores O-antigen expression on the bacterial cell surface. **B.** Analysis of phospholipid profiles from purified Δ*dolP* cell envelopes. Phospholipids were extracted by the Bligh-Dyer method from *E. coli* IM or OM samples purified by sucrose density gradient centrifugation. Phospholipids were visualized by staining with phosphomolybdic acid and charring after being separated by thin-layer chromatography with the following mobile phase: Chloroform:methanol:acetic acid (65:25:10). Phospholipid profiles were also analysed by LC/MS-MS following separation on the Luna C8(2) column under a THF/MeOH/H_2_O gradient. Phospholipid compositions are shown as sum for each of the four major classes observed: lyso-phophatidylethanolamines (LysoPE), phosphatidylethanolamines (PE), phosphatidylglycerols (PG) and cardiolipins (CL). Each data set is from three biological replicates generated from three separately purified membranes. Error bars represent ±S.D. **C.** PagP-mediated Lipid A palmitoylation assay. PagP transfers an acyl chain from surface exposed phospholipid to hexa-acylated Lipid A to form hepta-acylated Lipid A. [32P]-labelled Lipid A was purified from cells grown to mid-exponential phase in LB broth with aeration. Equal amounts of radioactive material (cpm/lane) was loaded on each spot and separated by thin-layer chromatography before quantification. As a positive control, cells were exposed to 25 mM EDTA for 10 min prior to Lipid A extraction in order to chelate Mg^2+^ ions and destabilize the LPS layer, leading to high levels of Lipid A palmitoylation. Hepta-acylated and hexa-acylated lipid A was quantified and hepta-acylated Lipid A represented as a percentage of total. Triplicate experiments were utilized to calculate averages and standard deviations with students t-tests used to assess significance. Student’s *t*-tests: NS* *P* > 0.1 compared with Parent EV. **D.***E. coli* BW25113 cells were grown overnight in LB (~16hrs) before being harvested by centrifugation and washed three times in PBS. Membrane fluidity was measured for each strain in triplicate and error bars represent standard deviation. Membrane fluidity is expressed as relative to *E. coli* BW25113 parent cells (WT).

**Fig. S11.**
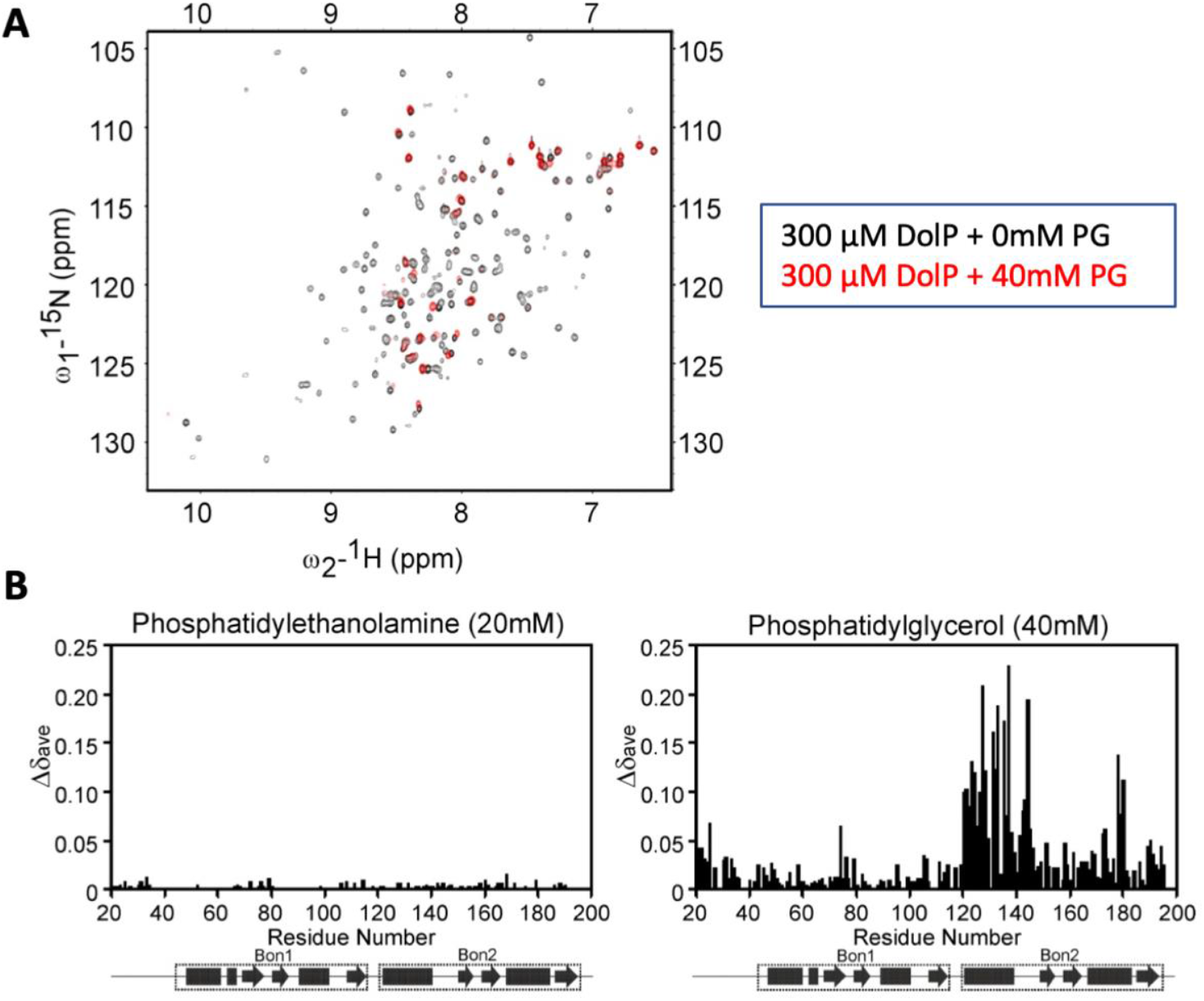
**A.** ^1^H,^15^N HSQC spectra of ^15^N-DolP (300 μM) in the presence (red) and absence (black) of 40 mM 1,2-dihexanoyl-sn-glycero-3-phospho-(1’-rac-glycerol) (DHPG) highlighting the large chemical shift perturbations observed on DHPG binding. **B.** Histograms showing the normalised CSP values observed in ^15^N labelled DolP (300 μM) amide signals in the presence of 5 mM cardiolipin, 20 mM 1,2,-dihexanoyl-sn-glycero-3-phosphethanolamine and 20 and 40 mM 1,2-dihexanoyl-sn-glycero-3-phospho-(1’-rac-glycerol).

**Fig. S12.**
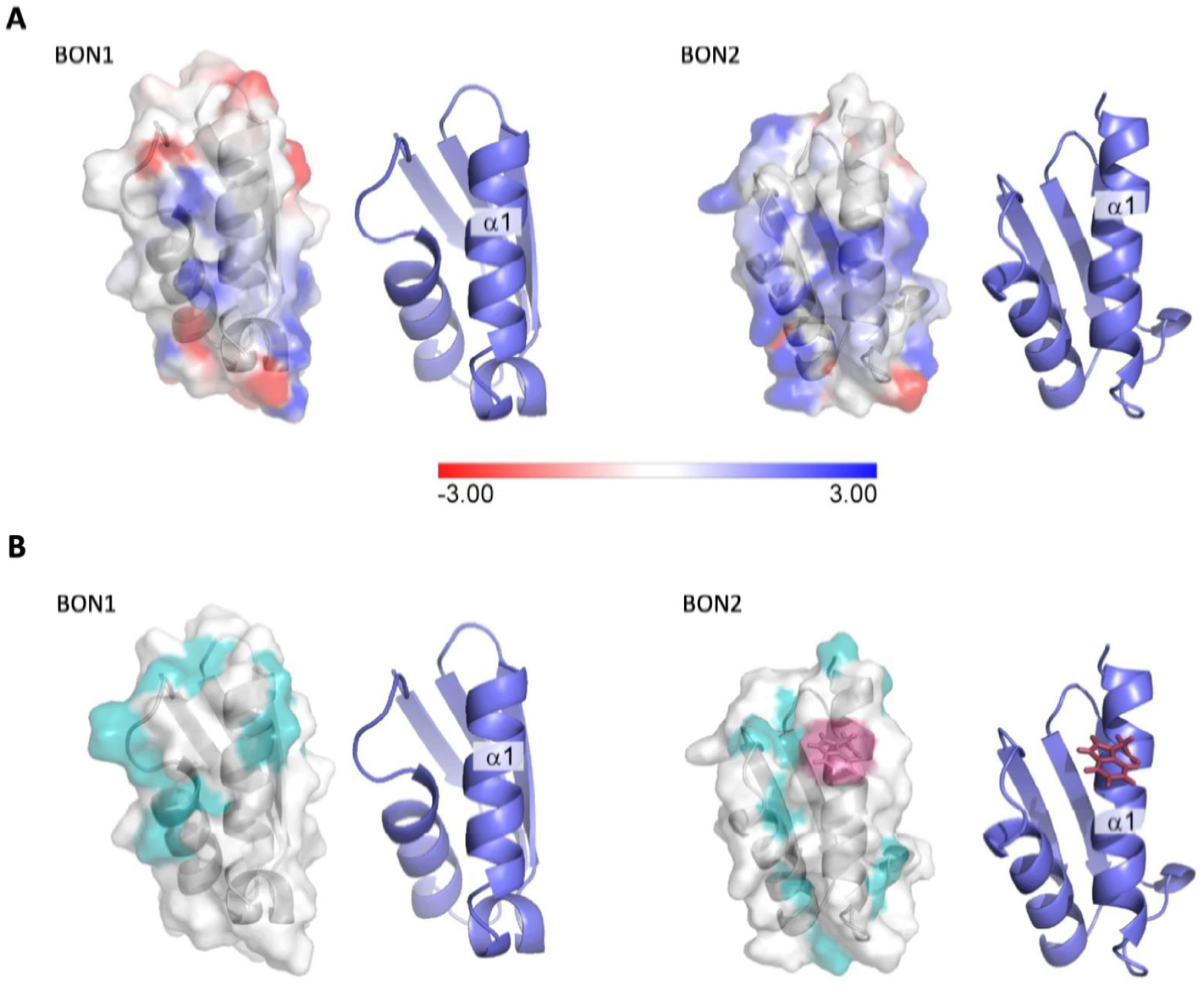
Electrostatic analysis of DolP. **A.** Electrostatic surface map of DolP BON domains 1 and 2 calculated using DelPhi^84^ at a pH of 6 and 0.05M ionic strength (which approximates the experimental conditions). The −3kT/e surface is shown in red and the +3kT/e surface is shown in blue. A formal charge library was used, with a dielectric of 2 assigned to the protein interior and a dielectric of 80 assigned to the exterior. Cartoon representations of the BON structures are shown to the right of each surface to more clearly highlight the orientations of the protein. The BON1:α1 and BON2:α1 helices show clear differences, with BON1:α1 being predominantly neutral with an electronegative patch towards its N-terminus, whilst BON2:α2 shows no electronegatively at all, but rather has a large electropositive patch towards the centre of this helix presumably explaining its specificity for the electropositive surface of phosphatidylglycerol. **B.** Hydrophobic surface map of DolP BON domains 1 and 2, hydrophobic residues (A, G, V, I, L, F, M) are shown in cyan, W127 (Red) is shown exposed on the surface of the BON2:α1 helix. Cartoon representations of the BON structures are shown to the right of each surface to more clearly highlight the orientations of the protein.

**Fig. S13.**
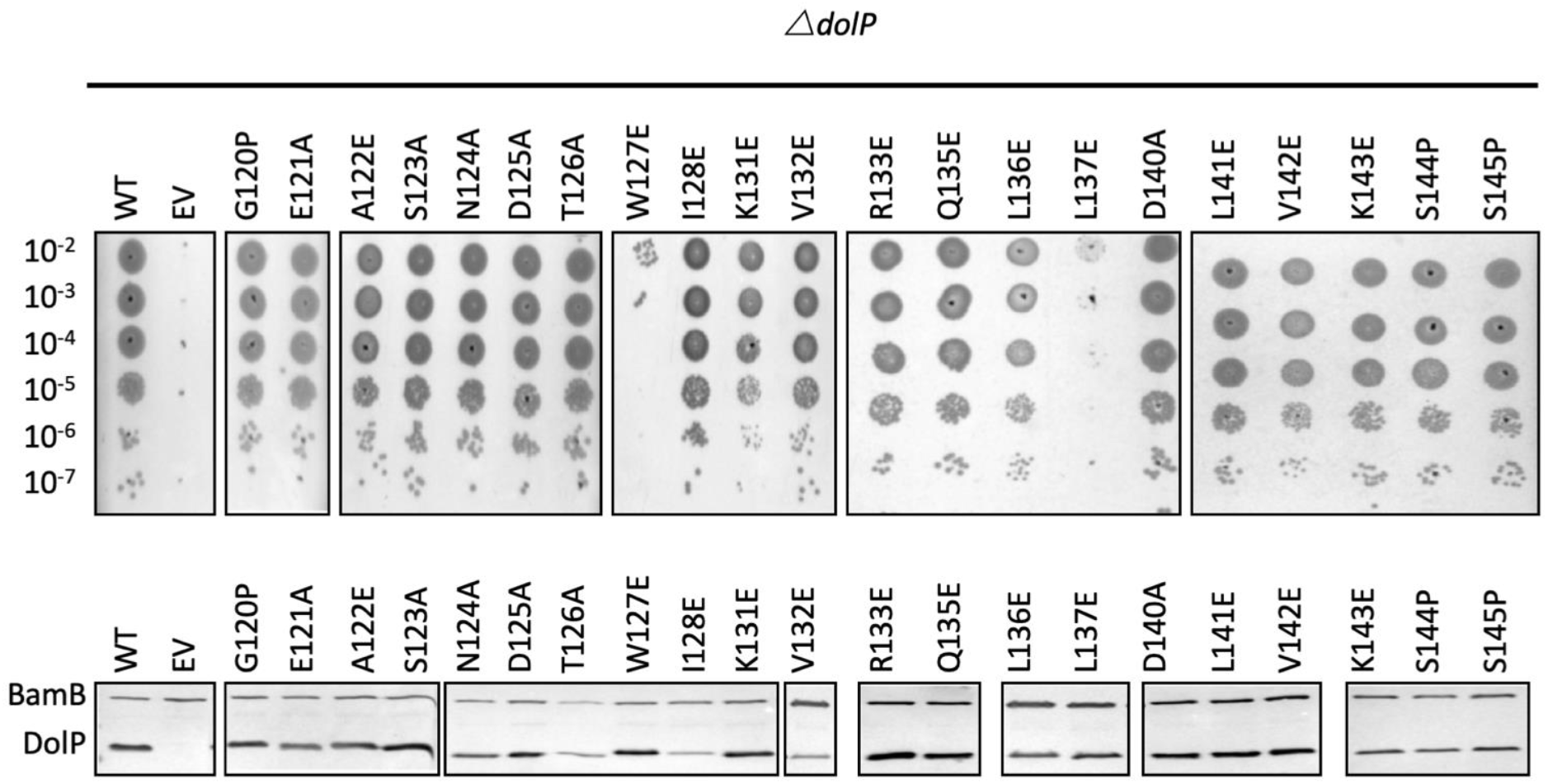
Analysis of DolP mutants. **A.***E. coli* BW25113 *ΔdolP* mutants were complemented with plasmids expressing a wild-type copy of DolP or a mutant version. Each strain was serially diluted and plated on LB-agar containing either vancomycin (100 μg/ml) or SDS (4.8% wt/vol) and growth was observed after overnight incubation. The W127E and L137E mutants failed to grow. **B.** Western immunoblotting of whole cell lysates derived from overnight cultures of mutants highlighted in the top panel. Blots were probed with antibodies to the outer membrane lipoprotein BamB and to DolP.

**Fig. S14.**
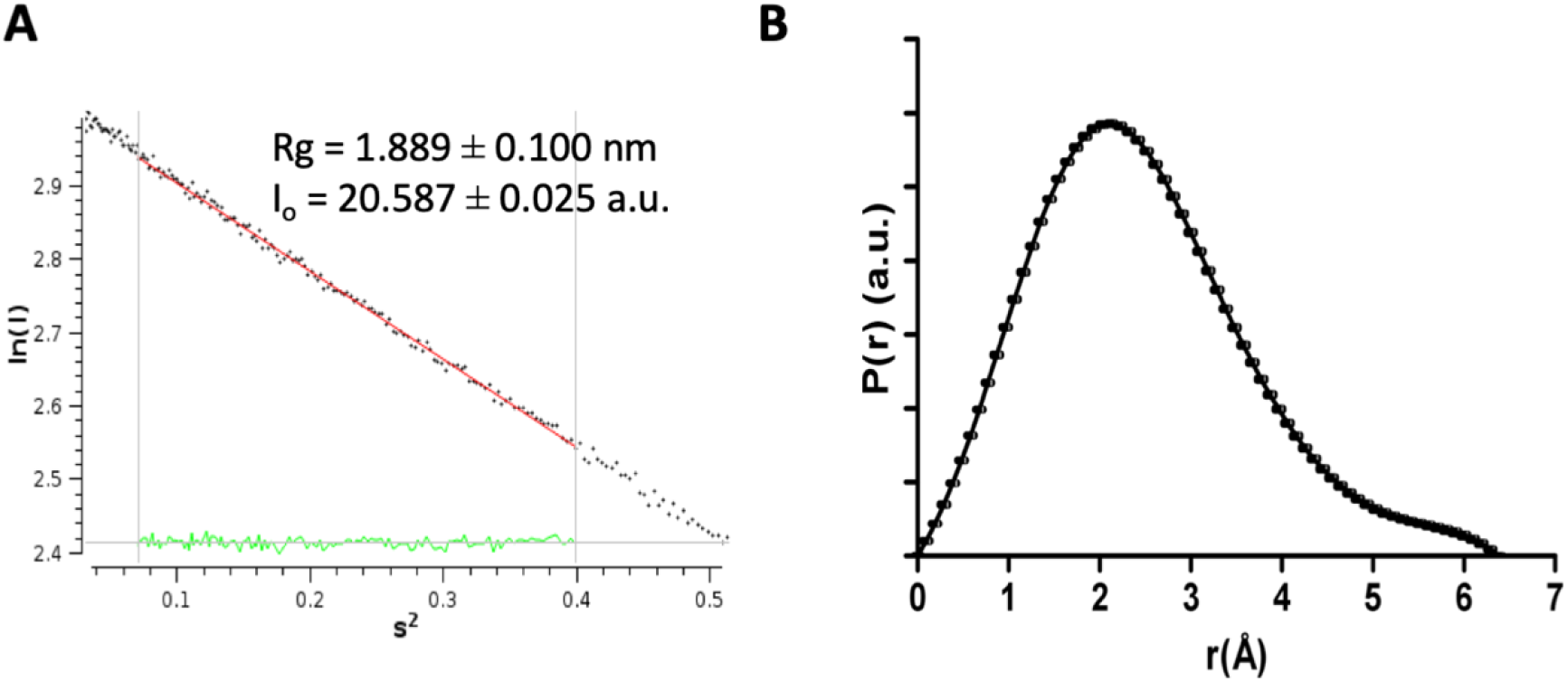
**A.** The linear region of the Guinier plot measured from the raw SAXS data for DolP. Values for Rg and I(0) are shown calculated using AutoRG in program Primus. **B.** Pair-wise distance distribution P(r), calculated from the scattering curve of DolP, calculated using gnom arbitrary units (a.u.).

**Table S1.**
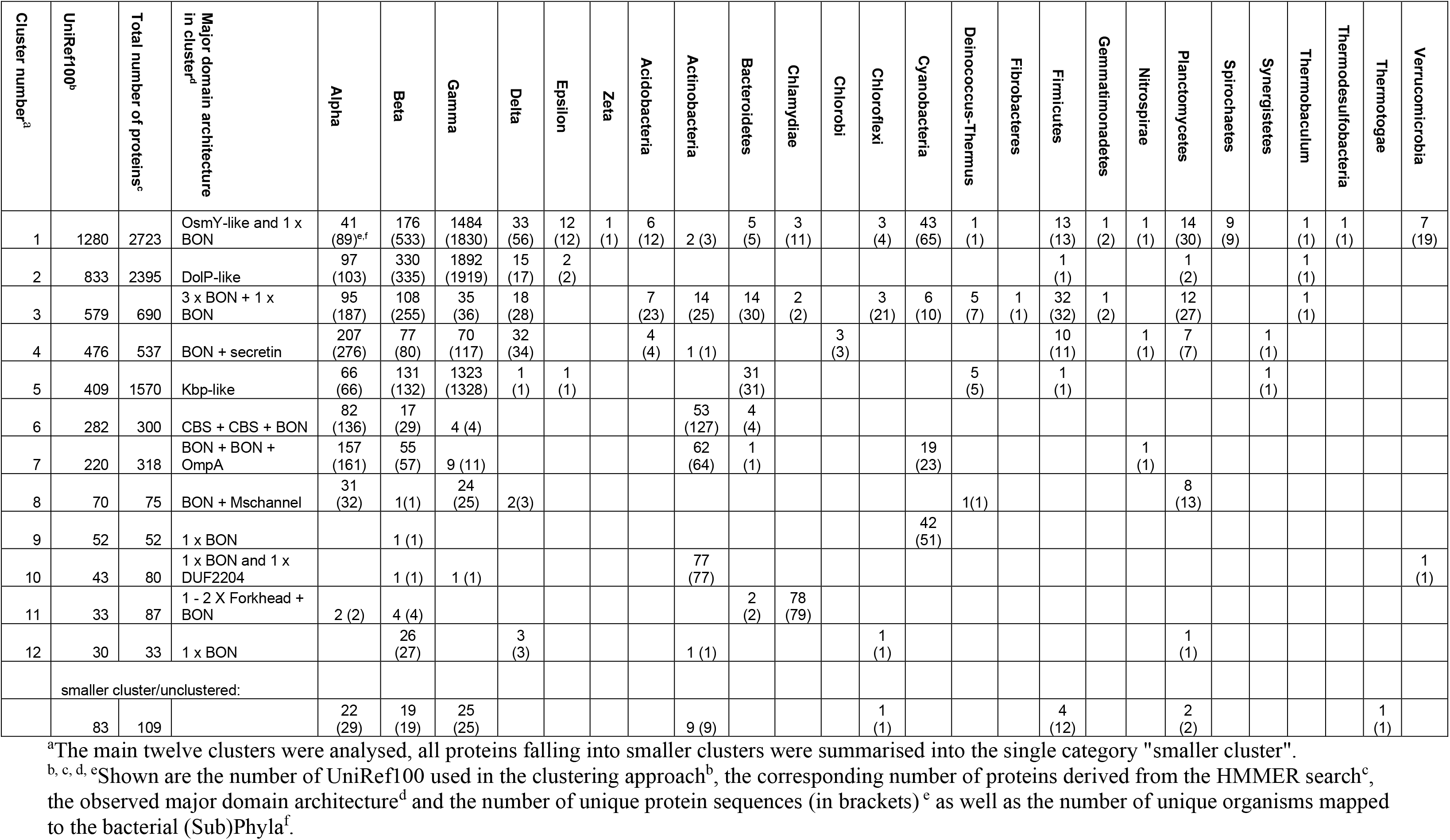
Taxonomic distribution of BON family domain architectures.

**Table S3.**
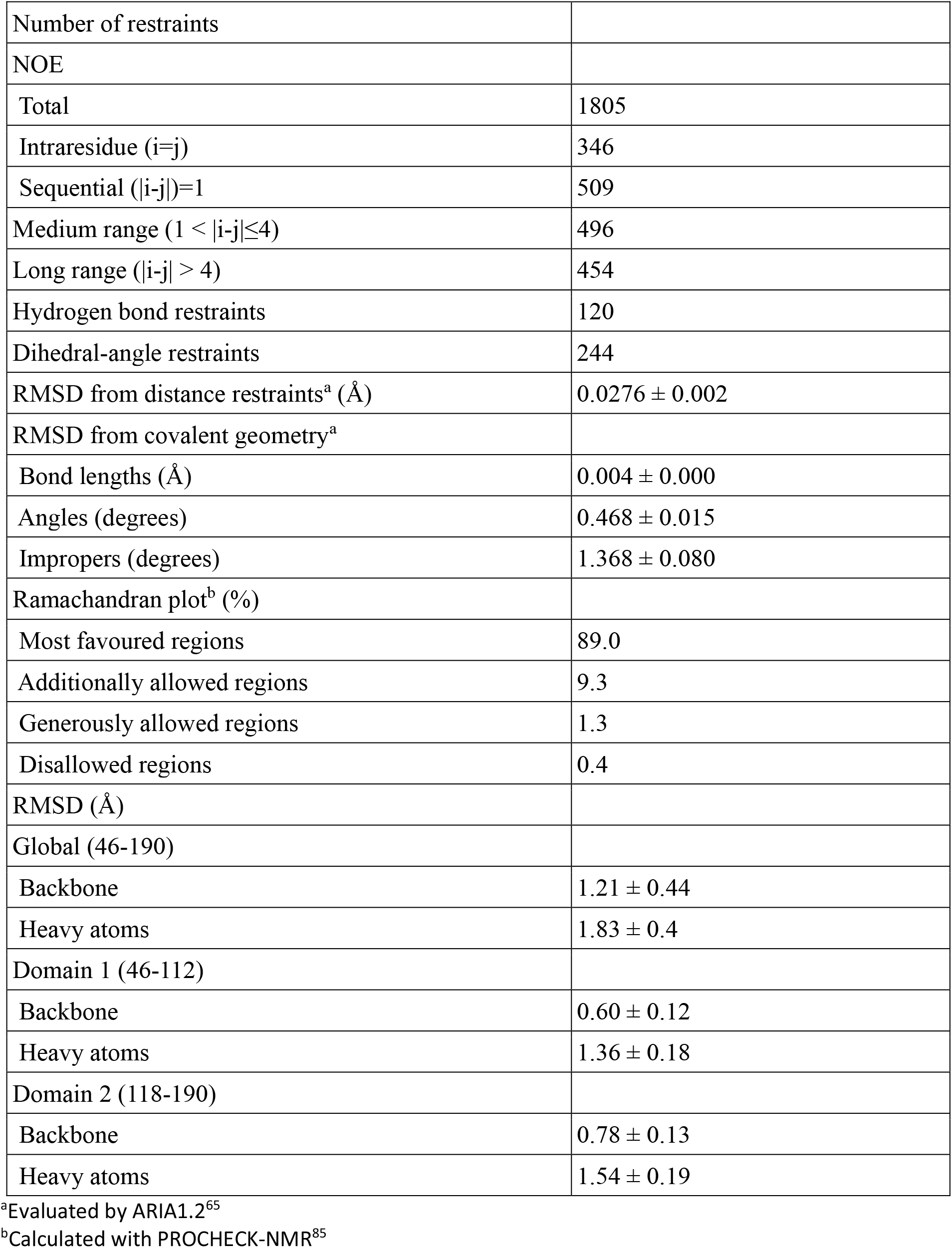
Structural statistics of the ensemble of 20 DolP solution structures

|  |  |
| --- | --- |
| Number of restraints |  |
| NOE |  |
| Total | 1805 |
| Intraresidue ($i=j$) | 346 |
| Sequential ($\vert i-j\vert =1$) | 509 |
| Medium range ($1 < \vert i-j\vert \leq 4$) | 496 |
| Long range ($\vert i-j\vert > 4$) | 454 |
| Hydrogen bond restraints | 120 |
| Dihedral-angle restraints | 244 |
| RMSD from distance restraints<sup>a</sup> (Å) | $0.0276 \pm 0.002$ |
| RMSD from covalent geometry<sup>a</sup> |  |
| Bond lengths (Å) | $0.004 \pm 0.000$ |
| Angles (degrees) | $0.468 \pm 0.015$ |
| Impropers (degrees) | $1.368 \pm 0.080$ |
| Ramachandran plot<sup>b</sup> (%) |  |
| Most favoured regions | 89.0 |
| Additionally allowed regions | 9.3 |
| Generously allowed regions | 1.3 |
| Disallowed regions | 0.4 |
| RMSD (Å) |  |
| Global (46-190) |  |
| Backbone | $1.21 \pm 0.44$ |
| Heavy atoms | $1.83 \pm 0.4$ |
| Domain 1 (46-112) |  |
| Backbone | $0.60 \pm 0.12$ |
| Heavy atoms | $1.36 \pm 0.18$ |
| Domain 2 (118-190) |  |
| Backbone | $0.78 \pm 0.13$ |
| Heavy atoms | $1.54 \pm 0.19$ |

<sup>a</sup>Evaluated by ARIA1.2<sup>65</sup>
<sup>b</sup>Calculated with PROCHECK-NMR<sup>85</sup>

**Table S5.** Accession numbers for the sequences used for CLANS clustering shown in Fig. 1

| Organism | OsmY | DolP | Kbp |
| --- | --- | --- | --- |
| *Escherichia coli* K12 | P0AFH8 | P64596 | P0ADE6 |
| *Klebsiella pneumoniae* MGH 78578 | A6THZ1 | A6TEG9 | A6T985 |
| *Enterobacter cloacae* ENHKU01 | J7G7C8 | J7GHD1 | J7GFT3 |
| *Salmonella enterica* Typhimurium | Q7CP68 | Q7CPQ6 | Q8ZML9 |
| *Erwinia billingiae* Eb661 | D8MMS8 | D8MME2 | D8MNV6 |
| *Serratia proteamaculans* 568 | A8G9G9 | A8GJZ3 | A8GFP7 |
| *Cronobacter sakazakii* ATCC BAA-894 | A7MGB6 | A7MIQ1 | A7MEA9 |
| *Pantoea* sp. Sc1 | H8DPK0 | H8DQ90 | H8DIH9 |
| *Hafnia alvei* ATCC 51873 | G9Y3J7 | G9Y4J4 | G9YAM4 |
| *Citrobacter rodentium* ICC168 | D2TRY4 | D2TQ24 | D2TM58 |
| *Shigella flexneri* 1235-66 | I6F1Q5 | I6GLP1 | I6HD15 |
| *Yersinia enterocolitica* 8081 | A1JJ93 | A1JR75 |  |
| *Yersinia pestis* KIM10+ | Q7CG58 | Q8D1R6 |  |
| *Dickeya dadantii* 3937 | E0SJX0 | E0SHF6 |  |

**Table S6.** HADDOCK docking statistics for ensemble 20 lowest energy DolP-DPC micelle solution structures calculated

|  |  |
| --- | --- |
| Experimental parameters<sup>a</sup> |  |
| Ambiguous distance restraints | 19 including NH of I20, G120-T130, V132-Q135, T138, S139 and NHε of W127 |
| Number of flexible residues<sup>b</sup> | 50 (I20-V45 (flexible linker as ascertained by NMR), A74, G120-I128, K131-R133, Q135-L137, V142-S145, I173, S178-V180) |
| Atomic pairwise RMSD (Å) |  |
| All backbone |  |
| Flexible interface backbone |  |
| Intermolecular energies (kcal.mol<sup>-1</sup>) |  |
| E<sub>vdw</sub> | -100.81 ± 7.74 |
| E<sub>elec</sub> | -231.67 ± 64.14 |
| E<sub>restraints</sub> | 22.30 ± 4.29 |
| Buried surface area (Å<sup>2</sup>) | 2186.78 ± 133.277 |

<sup>a</sup>deduced from intensity reductions observed in presence of 5-doxl derivative
<sup>b</sup>according to their surface accessibility and the chemical shift perturbation in presence of DPC/CHAPS

## References

1 May, K. L. & Grabowicz, M. The bacterial outer membrane is an evolving antibiotic barrier. Proc Natl Acad Sci U S A 115, 8852–8854 (2018).

2 Konovalova, A., Kahne, D. E. & Silhavy, T. J. Outer Membrane Biogenesis. Annu Rev Microbiol 71, 539–556 (2017).

3 Leyton, D. L., Rossiter, A. E. & Henderson, I. R. From self sufficiency to dependence: mechanisms and factors important for autotransporter biogenesis. Nat Rev Microbiol 10, 213–225 (2012).

4 Babu, M. M. et al. A database of bacterial lipoproteins (DOLOP) with functional assignments to predicted lipoproteins. J Bacteriol 188, 2761–2773 (2006).

5 Egan, A. J. F. Bacterial outer membrane constriction. Mol Microbiol 107, 676–687 (2018).

6 Ekiert, D. C. et al. Architectures of Lipid Transport Systems for the Bacterial Outer Membrane. Cell 169, 273–285.e217 (2017).

7 Stubenrauch, C. J. & Lithgow, T. The TAM: A Translocation and Assembly Module of the beta-Barrel Assembly Machinery in Bacterial Outer Membranes. EcoSal Plus 8(2019).

8 Gray, A. N. et al. Coordination of peptidoglycan synthesis and outer membrane constriction during *Escherichia coli* cell division. eLife 4(2015).

9 Goodall, E. C. A. et al. The Essential Genome of *Escherichia coli* K-12. mBio 9(2018).

10 Morris, F. C. et al. YraP Contributes to Cell Envelope Integrity and Virulence of S*almonella enterica* Serovar Typhimurium. Infect Immun 86(2018).

11 Bos, M. P., Grijpstra, J., Tommassen-van Boxtel, R. & Tommassen, J. Involvement of *Neisseria meningitidis* lipoprotein GNA2091 in the assembly of a subset of outer membrane proteins. J Biol Chem 289, 15602–15610 (2014).

12 Tsang, M. J., Yakhnina, A. A. & Bernhardt, T. G. NlpD links cell wall remodeling and outer membrane invagination during cytokinesis in *Escherichia coli*. PLoS Genet 13, e1006888 (2017).

13 Carlson, M. L. et al. Profiling the *Escherichia coli* membrane protein interactome captured in Peptidisc libraries. eLife 8(2019).

14 Babu, M. et al. Global landscape of cell envelope protein complexes in *Escherichia coli*. Nat Biotechnol 36, 103–112 (2018).

15 Typas, A. et al. Regulation of peptidoglycan synthesis by outer-membrane proteins. Cell 143, 1097–1109 (2010).

16 Ishida, T. et al. DiaA, a novel DnaA-binding protein, ensures the timely initiation of *Escherichia coli* chromosome replication. J Biol Chem 279, 45546–45555 (2004).

17 Dartigalongue, C., Missiakas, D. & Raina, S. Characterization of the *Escherichia coli* sigma E regulon. J Biol Chem 276, 20866–20875 (2001).

18 Yeats, C. & Bateman, A. The BON domain: a putative membrane-binding domain. Trends Biochem Sci 28, 352–355 (2003).

19 Cowles, C. E., Li, Y., Semmelhack, M. F., Cristea, I. M. & Silhavy, T. J. The free and bound forms of Lpp occupy distinct subcellular locations in *Escherichia coli*. Mol Microbiol 79, 1168–1181 (2011).

20 Webb, C. T. et al. Dynamic association of BAM complex modules includes surface exposure of the lipoprotein BamC. J Mol Biol 422, 545–555 (2012).

21 Storek, K. M. et al. The *Escherichia coli* β-Barrel Assembly Machinery Is Sensitized to Perturbations under High Membrane Fluidity. J Bacteriol 201(2019).

22 Knowles, T. J. et al. Structure and function of BamE within the outer membrane and the beta-barrel assembly machine. EMBO Rep 12, 123–128 (2011).

23 Dominguez, C., Boelens, R. & Bonvin, A. M. HADDOCK: a protein-protein docking approach based on biochemical or biophysical information. J Am Chem Soc 125, 1731–1737 (2003).

24 Oliver, P. M. et al. Localization of anionic phospholipids in *Escherichia coli* cells. J Bacteriol 196, 3386–3398 (2014).

25 Renner, L. D. & Weibel, D. B. Cardiolipin microdomains localize to negatively curved regions of *Escherichia coli* membranes. Proc Natl Acad Sci U S A 108, 6264–6269 (2011).

26 Mileykovskaya, E. & Dowhan, W. Visualization of phospholipid domains in *Escherichia coli* by using the cardiolipin-specific fluorescent dye 10-N-nonyl acridine orange. J Bacteriol 182, 1172–1175 (2000).

27 Laloux, G. & Jacobs-Wagner, C. How do bacteria localize proteins to the cell pole? J Cell Sci 127, 11–19 (2014).

28 Lugtenberg, E. J. & Peters, R. Distribution of lipids in cytoplasmic and outer membranes of *Escherichia coli* K12. Biochim Biophys Acta 441, 38–47 (1976).

29 Onufryk, C., Crouch, M. L., Fang, F. C. & Gross, C. A. Characterization of six lipoproteins in the sigmaE regulon. J Bacteriol 187, 4552–4561 (2005).

30 Yan, Z., Hussain, S., Wang, X., Bernstein, H. D. & Bardwell, J. C. A. Chaperone OsmY facilitates the biogenesis of a major family of autotransporters. Mol Microbiol 112, 1373–1387 (2019).

31 Typas, A. et al. High-throughput, quantitative analyses of genetic interactions in E. coli. Nat Methods 5, 781–787 (2008).

32 Wu, T. et al. Identification of a multicomponent complex required for outer membrane biogenesis in *Escherichia coli*. Cell 121, 235–245 (2005).

33 Hagan, C. L., Kim, S. & Kahne, D. Reconstitution of outer membrane protein assembly from purified components. Science 328, 890–892 (2010).

34 Gunasinghe, S. D. et al. The WD40 Protein BamB Mediates Coupling of BAM Complexes into Assembly Precincts in the Bacterial Outer Membrane. Cell Rep 23, 2782–2794 (2018).

35 Knowles, T. J., Scott-Tucker, A., Overduin, M. & Henderson, I. R. Membrane protein architects: the role of the BAM complex in outer membrane protein assembly. Nat Rev Microbiol 7, 206–214 (2009).

36 Rojas, E. R. et al. The outer membrane is an essential load-bearing element in Gram-negative bacteria. Nature 559, 617–621 (2018).

37 Giuliani, M. M. et al. A universal vaccine for serogroup B meningococcus. Proc Natl Acad Sci U S A 103, 10834–10839 (2006).

38 Pizza, M. et al. Identification of vaccine candidates against serogroup B meningococcus by whole-genome sequencing. Science 287, 1816–1820 (2000).

39 Punta, M. et al. The Pfam protein families database. Nucleic Acids Res 40, D290–301 (2012).

40 Finn, R. D., Clements, J. & Eddy, S. R. HMMER web server: interactive sequence similarity searching. Nucleic Acids Res 39, W29–37 (2011).

41 Enright, A. J., Van Dongen, S. & Ouzounis, C. A. An efficient algorithm for large-scale detection of protein families. Nucleic Acids Res 30, 1575–1584 (2002).

42 Hunter, S. et al. InterPro in 2011: new developments in the family and domain prediction database. Nucleic Acids Res 40, D306–312 (2012).

43 Frickey, T. & Lupas, A. CLANS: a Java application for visualizing protein families based on pairwise similarity. Bioinformatics 20, 3702–3704 (2004).

44 Baba, T. et al. Construction of *Escherichia coli* K-12 in-frame, single-gene knockout mutants: the Keio collection. Mol Syst Biol 2, 2006.0008 (2006).

45 Datsenko, K. A. & Wanner, B. L. One-step inactivation of chromosomal genes in Escherichia coli K-12 using PCR products. Proc Natl Acad Sci U S A 97, 6640–6645 (2000).

46 Kikuchi, S., Shibuya, I. & Matsumoto, K. Viability of an *Escherichia coli* pgsA null mutant lacking detectable phosphatidylglycerol and cardiolipin. J Bacteriol 182, 371–376 (2000).

47 Suzuki, M., Hara, H. & Matsumoto, K. Envelope disorder of *Escherichia coli* cells lacking phosphatidylglycerol. J Bacteriol 184, 5418–5425 (2002).

48 Browning, D. F. et al. Laboratory adapted *Escherichia coli* K-12 becomes a pathogen of *Caenorhabditis elegans* upon restoration of O antigen biosynthesis. Mol Microbiol 87, 939–950 (2013).

49 Browning, D. F. et al. Mutational and Topological Analysis of the *Escherichia coli* BamA Protein. PLoS One 8, e84512 (2013).

50 Isom, G. L. et al. MCE domain proteins: conserved inner membrane lipid-binding proteins required for outer membrane homeostasis. Sci Rep 7, 8608 (2017).

51 Dalebroux, Z. D. et al. Delivery of cardiolipins to the Salmonella outer membrane is necessary for survival within host tissues and virulence. Cell Host Microbe 17, 441–451 (2015).

52 Bligh, E. G. & Dyer, W. J. A rapid method of total lipid extraction and purification. Can J Biochem Physiol 37, 911–917 (1959).

53 Wu, H., Southam, A. D., Hines, A. & Viant, M. R. High-throughput tissue extraction protocol for NMR- and MS-based metabolomics. Anal Biochem 372, 204–212 (2008).

54 Teo, A. C. K. et al. Analysis of SMALP co-extracted phospholipids shows distinct membrane environments for three classes of bacterial membrane protein. Sci Rep 9, 1813 (2019).

55 Parham, N. J. et al. PicU, a second serine protease autotransporter of uropathogenic *Escherichia coli*. FEMS Microbiol Lett 230, 73–83 (2004).

56 Leyton, D. L. et al. Size and conformation limits to secretion of disulfide-bonded loops in autotransporter proteins. J Biol Chem 286, 42283–42291 (2011).

57 Selkrig, J. et al. Discovery of an archetypal protein transport system in bacterial outer membranes. Nat Struct Mol Biol 19, 506–510, S501 (2012).

58 Wisniewski, J. R., Zougman, A., Nagaraj, N. & Mann, M. Universal sample preparation method for proteome analysis. Nat Methods 6, 359–362 (2009).

59 Muhandiram, D. R. & Kay, L. E. Gradient-enhanced triple resonance three-dimensional NMR experiments with improved sensitivity. J Magn Reson B103, 203–216 (1994).

60 Delaglio, F. et al. NMRPipe: a multidimensional spectral processing system based on UNIX pipes. Journal of biomolecular NMR 6, 277–293 (1995).

61 Goddard, T. D. & Kneller, D. G. SPARKY 3. University of California San Francisco (2008).

62 Cornilescu, G., Delaglio, F. & Bax, A. Protein backbone angle restraints from searching a database for chemical shift and sequence homology. Journal of biomolecular NMR 13, 289–302 (1999).

63 Schanda, P., Kupce, E. & Brutscher, B. SOFAST-HMQC experiments for recording two-dimensional heteronuclear correlation spectra of proteins within a few seconds. Journal of biomolecular NMR 33, 199–211 (2005).

64 Guntert, P. Automated NMR structure calculation with CYANA. Methods in molecular biology 278, 353–378 (2004).

65 Linge, J. P., O’Donoghue, S. I. & Nilges, M. Automated assignment of ambiguous nuclear overhauser effects with ARIA. Methods in enzymology 339, 71–90 (2001).

66 Laskowski, R. A., Moss, D. S. & Thornton, J. M. Main-chain bond lengths and bond angles in protein structures. J Mol Biol 231, 1049–1067 (1993).

67 Koradi, R., Billeter, M. & Wuthrich, K. MOLMOL: a program for display and analysis of macromolecular structures. Journal of molecular graphics 14, 51–55, 29–32 (1996).

68 Dancea, F., Kami, K. & Overduin, M. Lipid interaction networks of peripheral membrane proteins revealed by data-driven micelle docking. Biophys J 94, 515–524 (2008).

69 Konarev, P. V. et al. PRIMUS - a Windows-PC based system for small-angle scattering data analysis. J Appl Cryst. 36, 1277–1282 (2003).

70 Guinier, A. La diffraction des rayons Xa ux tres petits angles; applicationa l’etude de phenomenes ultramicroscopiques. Ann. Phys. 12, 161–237 (1939).

71 D.I., S. Determination of the regularization parameter in indirect-transform methods using perceptual criteria. J. Appl. Crystallogr. 25, 495–503 (1992).

72 Franke, D. & Svergun, D. I. DAMMIF, a program for rapid ab-initio shape determination in small-angle scattering. J Appl Crystallogr 42, 342–346 (2009).

73 Volkov, V. V. & Svergun, D. I. Uniqueness of ab-initio shape determination in small-angle scattering. J. Appl. Cryst. 36, 860–864 (2003).

74 Bernado, P., Perez, Y., Svergun, D. I. & Pons, M. Structural characterization of the active and inactive states of Src kinase in solution by small-angle X-ray scattering. J Mol Biol 376, 492–505 (2008).

75 Petoukhov, M. V. & Svergun, D. I. Global rigid body modeling of macromolecular complexes against small-angle scattering data. Biophys J 89, 1237–1250 (2005).

76 Ducret, A., Quardokus, E. M. & Brun, Y. V. MicrobeJ, a tool for high throughput bacterial cell detection and quantitative analysis. Nat Microbiol 1, 16077 (2016).

77 Banzhaf, M. et al. Outer membrane lipoprotein NlpI scaffolds peptidoglycan hydrolases within multi-enzyme complexes in *Escherichia coli*. EMBO J 39, e102246 (2020).

78 Kritikos, G. et al. A tool named Iris for versatile high-throughput phenotyping in microorganisms. Nat Microbiol 2, 17014 (2017).

79 Chong, Z. S., Woo, W. F. & Chng, S. S. Osmoporin OmpC forms a complex with MlaA to maintain outer membrane lipid asymmetry in *Escherichia coli*. Mol Microbiol 98, 1133–1146 (2015).

80 Yim, H. H. & Villarejo, M. *osmY*, a new hyperosmotically inducible gene, encodes a periplasmic protein in *Escherichia coli*. J Bacteriol 174, 3637–3644 (1992).

81 Weber, A., Kogl, S. A. & Jung, K. Time-dependent proteome alterations under osmotic stress during aerobic and anaerobic growth in *Escherichia coli*. J Bacteriol 188, 7165–7175 (2006).

82 Ashraf, K. U. et al. The Potassium Binding Protein Kbp Is a Cytoplasmic Potassium Sensor. Structure 24, 741–749 (2016).

83 Lennon, C. W. et al. Folding Optimization In Vivo Uncovers New Chaperones. J Mol Biol 427, 2983–2994 (2015).

84 Li, L. et al. DelPhi: a comprehensive suite for DelPhi software and associated resources. BMC Biophys 4, 9 (2012).

85 Laskowski, R. A., MacArthur, M. W., Moss, D. S. & Thornton, J. M. PROCHECK: a program to check the stereochemical quality of protein structures. J Appl Crystallogr 26, 283–291 (1993).

